# Compartmentalization of mRNA and Translation by Desmosomes

**DOI:** 10.1101/2025.04.17.649424

**Authors:** Alec D’Alessandro, Kwabena Badu-Nkansah, Sophia Link, Daniel Hlavaty, Glenn Bjerke, Christopher V. Nicchitta, Rui Yi, Terry Lechler

## Abstract

Subcellular compartmentalization enables cells to concentrate and locally regulate functions. Here we find that translational machinery is compartmentalized to the cell cortex in epidermal cells. Furthermore, we observed broad enrichment of transcripts at the cortex, establishing a novel axis of mRNA organization in these cells. Mechanistically, we identified the desmosomal protein, desmoplakin, to be essential for cortical recruitment of both ribosomes and mRNAs, through distinct mechanisms. While mRNA localization is usually proposed to promote local translation, we found that most cortical transcripts were translationally repressed. These findings reveal spatial and transcript-dependent features of the cortical translatome. One mediator of this regulation is the RNA-induced Silencing Complex (RISC) which is also cortically enriched in a desmoplakin-dependent manner. Under homeostatic conditions, cortical RISC associates with transcripts for cell adhesion and the cytoskeleton. Upon wounding RISC is delocalized from the cortex and its associated transcripts become translationally upregulated. Together our data demonstrate a desmosome-dependent cortical compartmentalization of translation that dynamically responds to barrier perturbations, including wounding.

## INTRODUCTION

Cells must coordinate many biochemical functions within a shared cytoplasm. Much work has highlighted the organization of the cytoplasm at the level of organelles and their contacts, membrane-free condensates, and supramolecular complexes ^1,2^ . Compartmentalization of the cytoplasm has important roles in allowing discrete signaling, functions, and interactions to occur at specified cellular locales, potentially allowing cells to better insulate signals and or to concentrate pathways in a certain area ^3^ . One prominent example of this is the enrichment of many signaling pathways that occurs just beneath the plasma membrane ^4^.

In addition to signaling pathways, translation is compartmentalized in many cell types ^5^. In polarized cells, such as intestinal epithelia and neurons, ribosomes are enriched in apical areas and axons, respectively ^6,7^. In non-polarized cell, ribosomes are also enriched at distinct sites including intercalated discs in the heart and Z-bands in striated muscle ^8^. Neither the mechanisms nor the functions of these subcellular localizations is known. Currently, the best understood example of this is the sub-cellular localization of ribosomes to the endoplasmic reticulum (ER). This association of ribosomes with ER is largely dependent upon Sec61 ^9,10^, and is important for translation across the ER membrane. Therefore, there is a fundamental lack of understanding of how ribosomes and translation are localized to many discrete cellular locales.

In addition to ribosomes, many mRNAs show subcellular localization. Both zip code-based and more general mechanisms, such as mRNA stability, have been proposed to mediate mRNA localization ^11^. In only a select few cases is the machinery that anchors mRNAs at specific sites known ^12^. It has also generally been assumed, but not always tested, that the translation of transcripts is also enriched at their site of localization, i.e. that the purpose of transcript localization is translation/protein localization. However, subcellular compartmentalization could also allow differential regulation of transcripts at distinct sites.

Here we report that the cell cortex serves as a distinct compartment for translation in epidermal cells, with enrichment of ribosomes, a subset of mRNAs, and additional translational regulators including the RNA-induced silencing complex (RISC). Notably, localization of both transcripts and translational machinery depend upon desmosomes, stable cell-cell adhesions that are canonically known for providing mechanical resilience to the skin. Molecularly, the recruitment of ribosomes and mRNAs are mediated by distinct domains of the core desmosomal protein, desmoplakin. Notably, many of the transcripts enriched at the cell cortex are translationally repressed. In contrast, there is active cortical translation of mRNAs encoding cell adhesion and cytoskeletal proteins. These data demonstrate local regulation of translation that is transcript dependent. We find that one mechanism of local translational repression is provided by the RISC complex, which localizes to the cortex where it binds to transcripts enriched in cell adhesion genes. Wounding induces a decrease in cortical RISC and an increase in the translation of many RISC-associated transcripts. Remarkably, cell adhesion and cytoskeletal proteins are among the most highly translationally increased in response to wounding, suggesting that disrupting cell adhesion results in rapid translational changes to restore epithelial integrity.

## RESULTS

### Ribosomes localize to the cortex in a desmoplakin-dependent manner

Desmosomes are cell adhesions whose canonical role is anchoring the intermediate filament (IF) cytoskeleton ^13–15^. However, recent data demonstrate that they are multi-functional organizers of the cytoplasm, capable of regulating actin, microtubules, and the endoplasmic reticulum ^16–20^. We recently used a proximity-biotinylation approach (BioID) to elucidate the desmosome proteome (Fig. 1A)^21^ . Gene ontology analysis (by cellular component) of desmosome-associated proteins revealed cell adhesion as the most highly enriched term. Surprisingly, translation, RNA-binding, and translational regulation were also highly significantly enriched GO terms (Fig. 1B). Within these categories, ribosomes, translation initiation factors, elongation factors, and tRNA synthetases were all enriched in the desmosome-associated fraction, suggesting broad localization of translational machinery (Fig. 1C and Supplemental Table 1). Structural mapping of the BioID-identified ribosomal proteins onto the ribosome revealed that most surfaces of this macromolecule were efficiently biotinylated (Fig. 1D). We also found almost complete coverage of the RNA-induced silencing complex (RISC) which plays roles in the stability and translational regulation of mRNAs ^22^. To validate these findings, we started with an approach orthogonal to the biochemical screen, using immunofluorescence to characterize the localization of individual proteins. We co-stained for the core desmosomal protein, desmoplakin, and ribosomal proteins (RPS3, small subunit; and RPL10A, large subunit) in wild-type mouse keratinocytes. Both small subunit and large subunit proteins were clearly enriched at the cell cortex in the vicinity of the desmosomes (Fig. 2A,D), which was confirmed using line scan analysis to measure signal across the cell cortex (Fig. 2B,C,E,F). Notably, RPL10A was not identified in the BioID screen (peptides were not recovered in either the experimental or control conditions), but was clearly cortically localized. These data suggest that most/all components of the ribosome are cortically localized, with a subset of them not being well-labeled by the BioID approach. There was also a cortical pool of RPL10A and RPS3 in HaCaT cells, a human epidermal cell line, demonstrating conservation of ribosome localization across species (Fig. S1A). To control for potential antibody artifacts, we examined primary mouse keratinocytes expressing an EGFP-tagged RPL10A protein (Fig. S1B). This tagged protein showed a strong cortical enrichment as compared to the cytoplasmic EGFP control. Ribosomes were also cortically enriched in intact mouse skin, most clearly in the terminally differentiated granular cells of this tissue (Fig. S1C). Finally, an RNA binding protein identified in our screen, PolyA binding protein C (PABPC), was also found to be cortically enriched (Fig. S1D).

**Figure 1:**
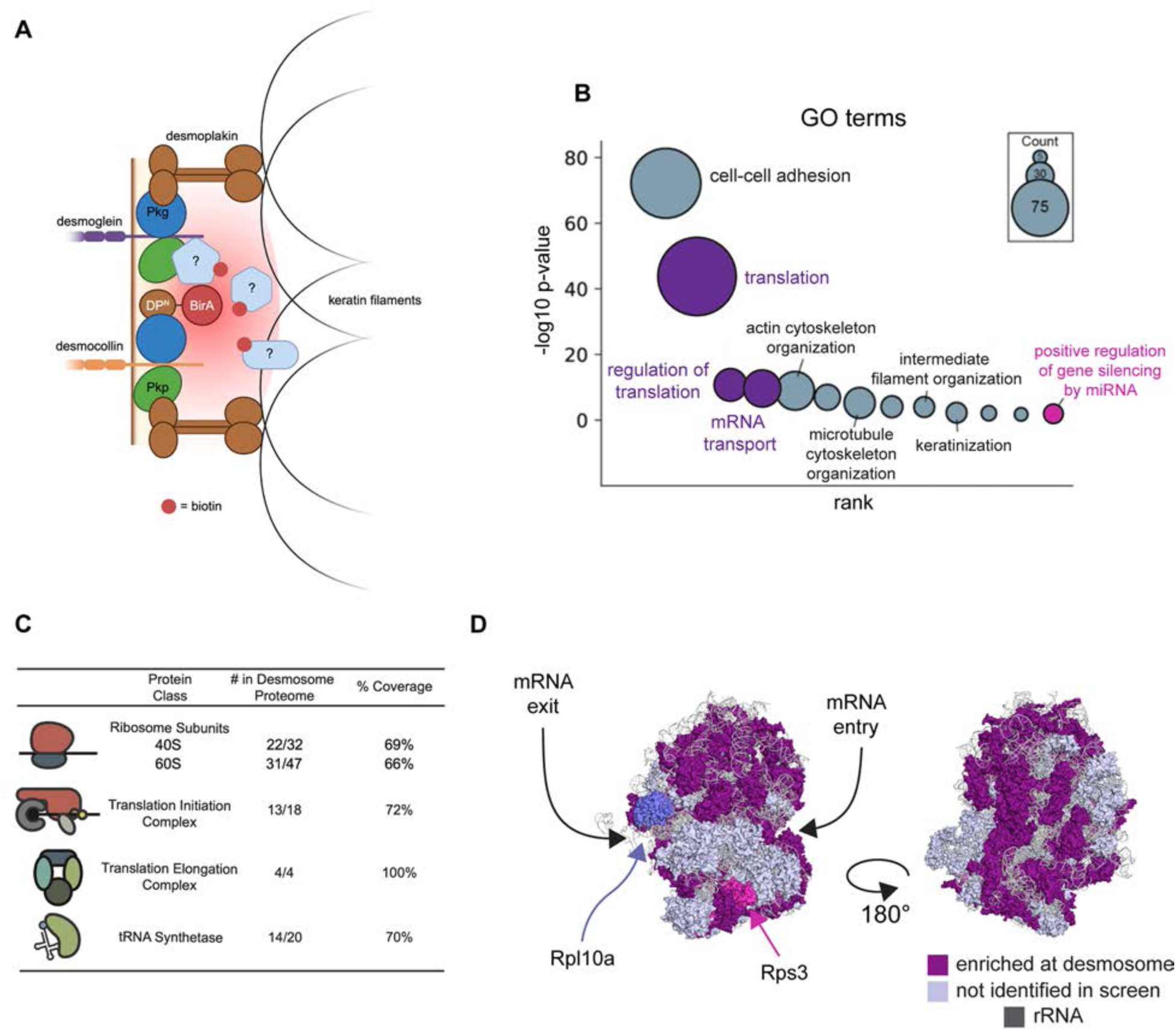
Translation machinery is recovered in desmosome BioID. (A) Schematic of BioID screen to identify proteins enriched at desmosomes. (B) GO terms of proteins identified as enriched at desmosomes listed by rank, significance, and count. (C) Breakdown of translation machinery identified as enriched at desmosomes. (D) Representation of human 80S ribosome (PDB 4UG0) with color-scheme to show positions of ribosomal proteins identified as enriched at desmosomes.

**Figure 2:**
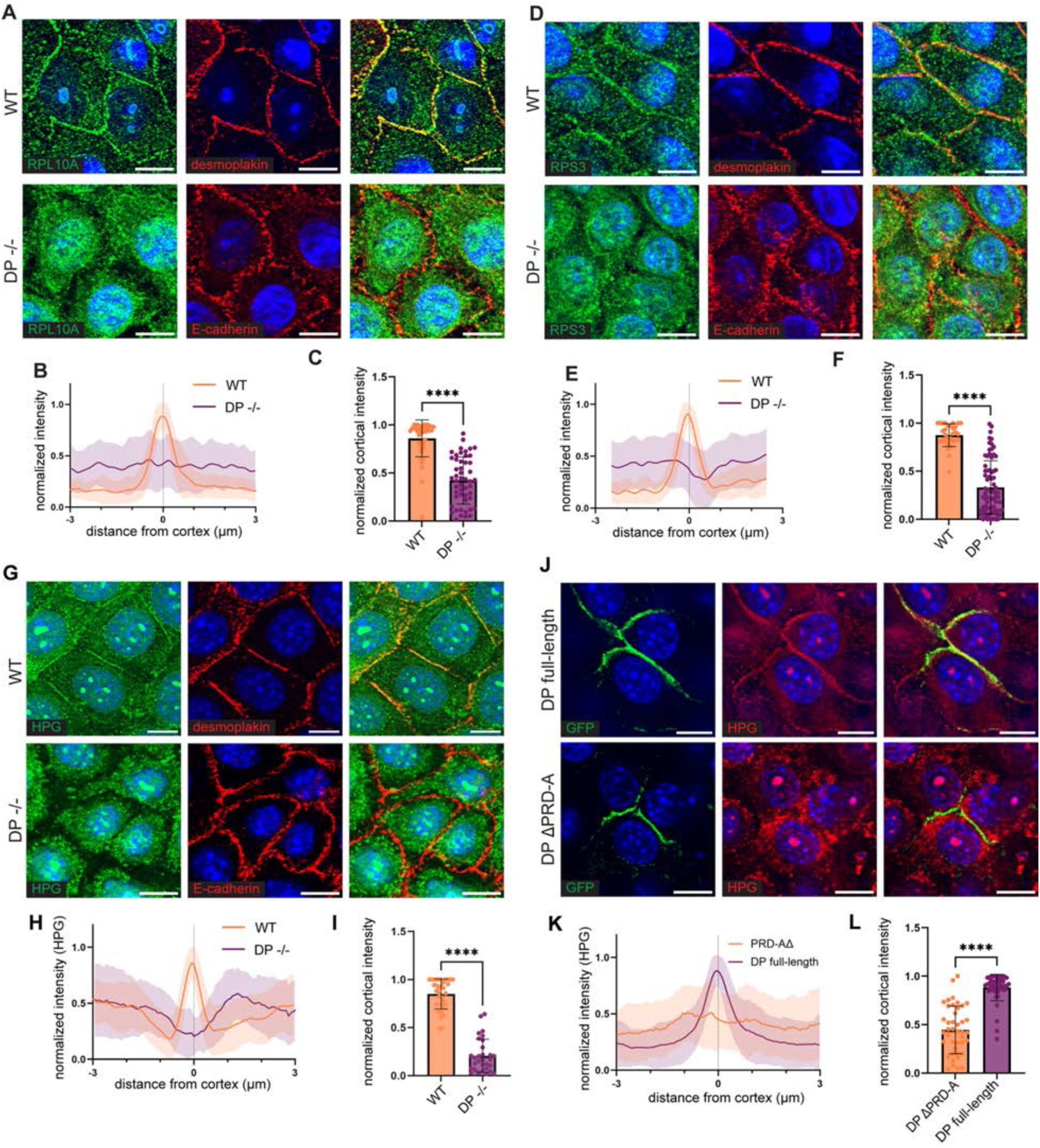
Ribosomes and translation are localized to the cell cortex in a desmoplakin-dependent manner. (A) Immunofluorescence of RPL10A and desmoplakin or E-cadherin in wild-type or desmoplakin-null mouse keratinocytes (scale bar = 10 µm). (B) Line scans of RPL10A fluorescence intensity measured across the cell cortex. Shaded area represents the standard deviation (WT n = 42 cells, DP-/- n = 50 cells; 3 biological replicates) (C) Relative intensity of RPL10A fluorescence at the center of the cell-cell boundary (WT n = 42, DP-/- n = 50; 3 biological replicates; p < 0.0001). (D) Immunofluorescence of RPS3 and desmoplakin or E-cadherin in wild-type or desmoplakin-null mouse keratinocytes (scale bar = 10 µm). (E) Line scans of RPS3 fluorescence intensity measured across the cell cortex. Shaded area represents the standard deviation (WT n = 30 cells, DP-/- n = 75 cells; 3 biological replicates). (F) Relative intensity of RPS3 fluorescence at the center of the cell-cell boundary (WT n = 30 cells, DP-/- n = 75 cells; 3 biological replicates; p < 0.0001). (G) Click-chemistry labeling of L-homopropargylglycine (HPG; 3 min pulse) and immunofluorescence of desmoplakin or E-cadherin in wild-type or desmoplakin-null mouse keratinocytes (scale bar = 10 µm). (H) Line scans of HPG fluorescence intensity measured across the cell cortex of wild-type and desmoplakin-null keratinocytes. Shaded area represents the standard deviation (WT n = 33 cells, DP-/- n = 30 cells; 3 biological replicates). (I) Relative intensity of HPG fluorescence at the center of the cell-cell boundary (WT n = 33 cells, DP-/- n = 30 cells; 3 biological replicates; p < 0.0001). (J) Desmoplakin-null keratinocytes transfected with plasmids expressing GFP-tagged full-length or ΔPRD-A desmoplakin constructs pulsed with HPG (3 min). (K) Line scans of HPG fluorescence intensity measured across the cell cortex of transfected keratinocytes. Shaded area represents the standard deviation (DP full-length n = 45 cells, ΔPRD-A n = 42 cells; 3 biological replicates). (L) Relative intensity of HPG fluorescence at the center of the cell-cell boundary (DP full-length n = 45 cells, ΔPRD-A n = 42 cells; 3 biological replicates; p < 0.0001).

Given that we identified ribosomes in a desmosome proteomics screen, we next tested whether desmosomes were important for ribosomal cortical localization. In both cells and intact epidermal tissue null for desmoplakin, ribosomes were no longer localized to the cell cortex (Fig. 2, Fig. S1C). This was also true for PABPC (Fig. S1D). This effect was specific to desmosomes and not a general result of disrupting cell-cell adhesion as loss of p120-catenin, a stabilizer of adherens junctions, did not alter the cortical localization of ribosomes (compare Fig. S1E to 2A) ^23,24^.

Desmoplakin is a multi-functional protein that clusters desmosomes and directly links them to keratin filaments. To determine whether cortical recruitment of ribosomes is linked to keratin binding/organization at desmosomes, we examined ribosome localization in cells null for type II keratins which are necessary for filament formation ^25^. Ribosomes localized to the cortex in these cells demonstrating that keratins are dispensable for cortical ribosome localization (Fig. S1F). We also tested the roles of the actin and microtubule cytoskeletons, both of which are known to be organized by desmosomes ^17–19^. Neither treatment of cells with the actin depolymerizing drug, cytochalasin B, nor the microtubule depolymerizing drug, nocodazole, resulted in loss of the cortical localization of ribosomes, even after 1 hour of treatment (Fig. S1G), demonstrating that they are not required for maintenance of localization. In addition, active translation was not required for localization as neither cycloheximide nor puromycin treatment affected cortical ribosome enrichment (Fig. S1G).

### Molecular mechanisms of cortical ribosome recruitment

To gain more insight into molecular requirements for cortical ribosome localization, we performed rescue experiments to understand which domains of desmoplakin are required for localization. The desmoplakin head domain mediates desmosome stability by binding desmosomal cadherins and cytoplasmic desmosome components, while its tail domain interacts with keratin filaments ^26–28^. Expression of full length desmoplakin rescued cortical ribosome localization in desmoplakin null cells (Fig. 3A-C). In contrast, desmoplakin lacking the tail domain (DPΛ1tail) was unable to do so. An internal deletion of the dimerizing coiled-coil region of desmoplakin, which preserves the IF-binding tail domain, resulted in a protein that was able to rescue ribosomal localization. Together, these data pointed to a role for desmoplakin’s tail domain in ribosome recruitment, demonstrating a function for this region beyond IF interaction.

**Figure 3:**
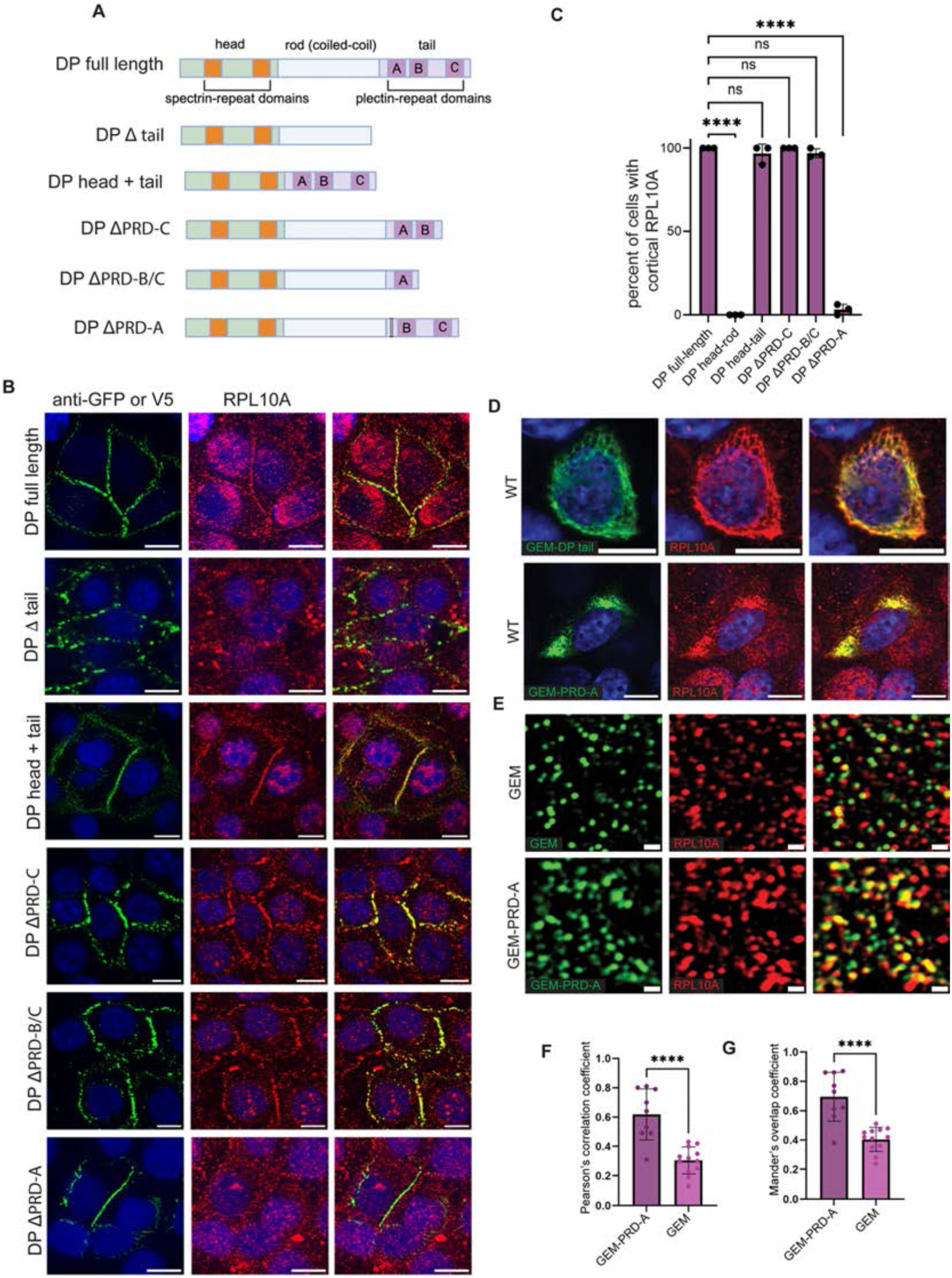
The PRD-A domain of desmoplakin is necessary and sufficient for ribosomal localization. (A) Overview of desmoplakin constructs tested for their ability to rescue ribosome and localization. (B) Desmoplakin-null cells transfected with various GFP or V5-tagged desmoplakin constructs and stained for GFP or V5 and RPL10A. (C) Percent of desmoplakin-null cells transfected with various desmoplakin constructs that show rescued cortical RPL10A immunofluorescence (n = >10 cells from 3 biological replicates; **** p < 0.0001). (D) Wild-type keratinocytes transfected with GEM-DP tail or GEM-PRD-A and co-stained for RPL10A (scale bar = 10 µm). (E) Lattice-SIM images of wild-type mouse keratinocytes transfected with GEM-PRD-A or control GEM co-stained for RPL10A (scale bar = 100 nm). (F) Pearson’s correlation coefficient (PCC) from lattice-SIM images of RPL10A and GEM-PRD-A or control GEM fluorescence in wild-type mouse keratinocytes (GEM-PRD-A n = 9 cells, GEM n = 12 cells; p < 0.0001). (G) Mander’s overlap coefficient (MOC) from lattice-SIM images of RPL10A and GEM-PRD-A or GEM fluorescence in wild-type mouse keratinocytes (GEM-PRD-A n = 9 cells, GEM n = 12 cells; p < 0.0001).

Desmoplakin’s tail domain contains three plectin-repeat domains (PRD), named PRD-A, B, and C. While PRD-B and C show robust IF interaction ability *in vitro* and *in vivo*, PRD-A is distinct in having low affinity for keratins ^29^. Deletion of either the PRD-C or both PRD-B and -C did not affect cortical ribosome recruitment (Fig. 3B,C), suggesting that this activity likely resides in PRD-A and/or adjacent sequences. We directly tested this by deleting the PRD-A domain in the context of the full-length protein and found that this construct was unable to rescue ribosome recruitment to the cortex (Fig. 3B,C). To ensure that this effect was not simply due to different levels of expression of the mutant construct, we also determined the ratio of RPL10A signal to EGFP fluorescence (Fig. S2A), which further demonstrated a necessity for the PRD-A domain in cortical ribosome recruitment. The PRD-A deletion construct was able to localize to the cortex and to rescue keratin organization, suggesting that its other major functions remain intact.

To investigate whether the desmoplakin tail domain was sufficient to interact with ribosomes, we fused the desmoplakin tail to a <u>g</u>enetically <u>e</u>ncoded <u>m</u>ultimeric protein (GEM) derived from the encapsulin protein of *Pyrococcus furiosus* ^30^ . This allows formation of approximately 40 nm structures, which, while still smaller than desmosomes, mimic the high local concentration and avidity found there. When expressed in wild-type mouse keratinocytes, this fusion protein decorated keratin filaments throughout the cell (Fig. 3D). Moreover, when staining for ribosomes, we observed a strong colocalization with the GEM-DP tail fusion protein, demonstrating that the tail is sufficient to reorganize ribosome localization. We next tested this construct in keratin null cells, to rule out possible effects of keratin binding. Under these conditions the GEM-DP tail tended to aggregate, and we again saw colocalization between the GEM-DP tail fusion protein and ribosomes (Fig. S2B). Colocalization with ribosomes was not seen when expressing the GEM protein alone, or with another GEM-fusion that also resulted in large aggregate formation (Fig. S2C). Further, a GEM-PRD-A fusion was sufficient to colocalize with ribosomes (Fig. 3E-G).

To further assess possible interactions between the PRD-A domain and ribosomes, we performed polysome analysis on cells expressing a cytoplasmic PRD-A. We found that this domain co-sedimented with ribosomal subunits, monosomes and polysomes (Fig S2D). Further, disrupting ribosomes or polysomes with either EDTA or RNase A caused loss of PRD-A from the heavy fractions (Fig. S2E,F), demonstrating that PRD-A sedimentation is dependent upon ribosomes. Together, these data demonstrate a molecular mechanism for cortical ribosome localization, mediated by the PRD-A domain of desmoplakin. To our knowledge, this is the first molecular mechanism for ribosome subcellular localization.

### Active cortical translation in keratinocytes

Given the localization of ribosomes at the cortex, we next tested whether active translation occurred at this cellular site. To observe translation, we first used the puromycin analog, O-propargyl-puromycin (OPP), which is incorporated onto the carboxyl terminus of nascent peptides and halts translation ^31^. OPP contains an alkyne group that allows for click-chemistry labeling with fluorophores. We pulsed wild-type mouse keratinocytes for 5 min with OPP, followed by rapid fixation, click-chemistry conjugation of fluorescent dyes, and counterstaining with desmoplakin antibodies. We observed that OPP signal was enriched at the cortex in the vicinity of desmosomes (Fig. S3A-C). This suggested that at least a portion of the ribosomes localized to the cortex were actively translating. Because puromycin can lead to peptide release from the ribosome, potentially causing artefactual results ^32,33^, we performed a similar assay using L-homopropargylglycine (HPG), a methionine analog that is incorporated into nascent peptides that can also be labelled via click-chemistry ^34^. HPG is non-chain terminating, so labeling should reflect translating peptides as well as recently released proteins. HPG incorporation was detected throughout the cell including colocalizing with desmoplakin at the cortex in both mouse and human keratinocytes (Fig. 2G-I; S3E,F). These localizations were also observed using SIM to achieve a higher resolution view (Fig. S3G,H). Cortical translation was lost in desmoplakin null cells, consistent with the loss of cortical ribosome localization (Fig. 2G-I;Fig. S3A-C). This effect was specific to desmosomes, as loss of p120-catenin did not result in loss of cortical translation (Fig. S3A,D).

To rule out possible cortical capture of nascent proteins, we turned to rescue experiments. Re-expression of desmoplakin in desmoplakin null cells restored the cortical HPG signal (Fig. 2J-L). Notably, cells expressing desmoplakin lacking the PRD-A domain showed no cortical HPG signal, despite the localization of this protein to the cortex (Fig. 2J, Fig. 3B). This is strong evidence that this domain is important for cortical translation, likely through recruitment of ribosomes, and largely rules out the possibility that nascent peptides are simply accumulating at the cortex after ribosomal release. Additionally, if the HPG assay was simply a readout for sites of nascent peptide accumulation rather than sites of translation, then recruitment of ribosomes to another site within the cell should not alter HPG labeling patterns. However, we find that recruitment of ribosomes to keratin filaments using the GEM-DP-tail construct (described above) alters the localization of HPG incorporation, co-localizing with the GEM construct and keratin filaments (Fig S3I,J). Together, these experiments validate that HPG signal reflects the site of translation, and that the cortical HPG signal observed is not simply captured nascent peptides that are released from ribosomes.

We next turned to polysome analysis to further interrogate the status of cortical ribosomes. To isolate ribosomes from the cortical compartment, we expressed the DP-BirA construct to biotinylate cortical proteins. The cells were then lysed and monosomes/polysomes were separated on a sucrose gradient (Fig. S4A). Fractions were incubated with streptavidin resin to isolate cortical ribosomes, yielding monosome and polysome fractions from the cytoplasm and cortex. Western blotting for RPL10A allowed us to calculate a relative ratio between monosomes and polysomes (Fig. S4B). For both the cytoplasm and the cortex, the majority of ribosomes were associated with polysomes. However, relative to the cytoplasm, the cortex had a higher polysome-to-monosome ratio (Fig. S4C). This provides an additional line of evidence that there is active translation at the cortex.

### A subset of mRNAs localizes to the cell cortex

The presence of active cortical translation raised the question of whether mRNAs at the cortex represented a similar or distinct composition as compared to the cytoplasm. To specifically mark and isolate cortical mRNAs we took advantage of our DP-BirA keratinocyte cell line. This allowed for biotinylation of cortical mRNA binding proteins (and potentially, direct labeling of mRNAs at low efficiency) and the isolation of mRNAs after streptavidin pulldown (Fig. 4A). Sequencing of cortical and cytoplasmic mRNAs revealed broad transcript compartmentalization in these cells. In total, 3615 mRNAs (25%) were enriched at the cortex, while 3477 mRNAs were depleted (24%) (Fig. 4B).

**Figure 4:**
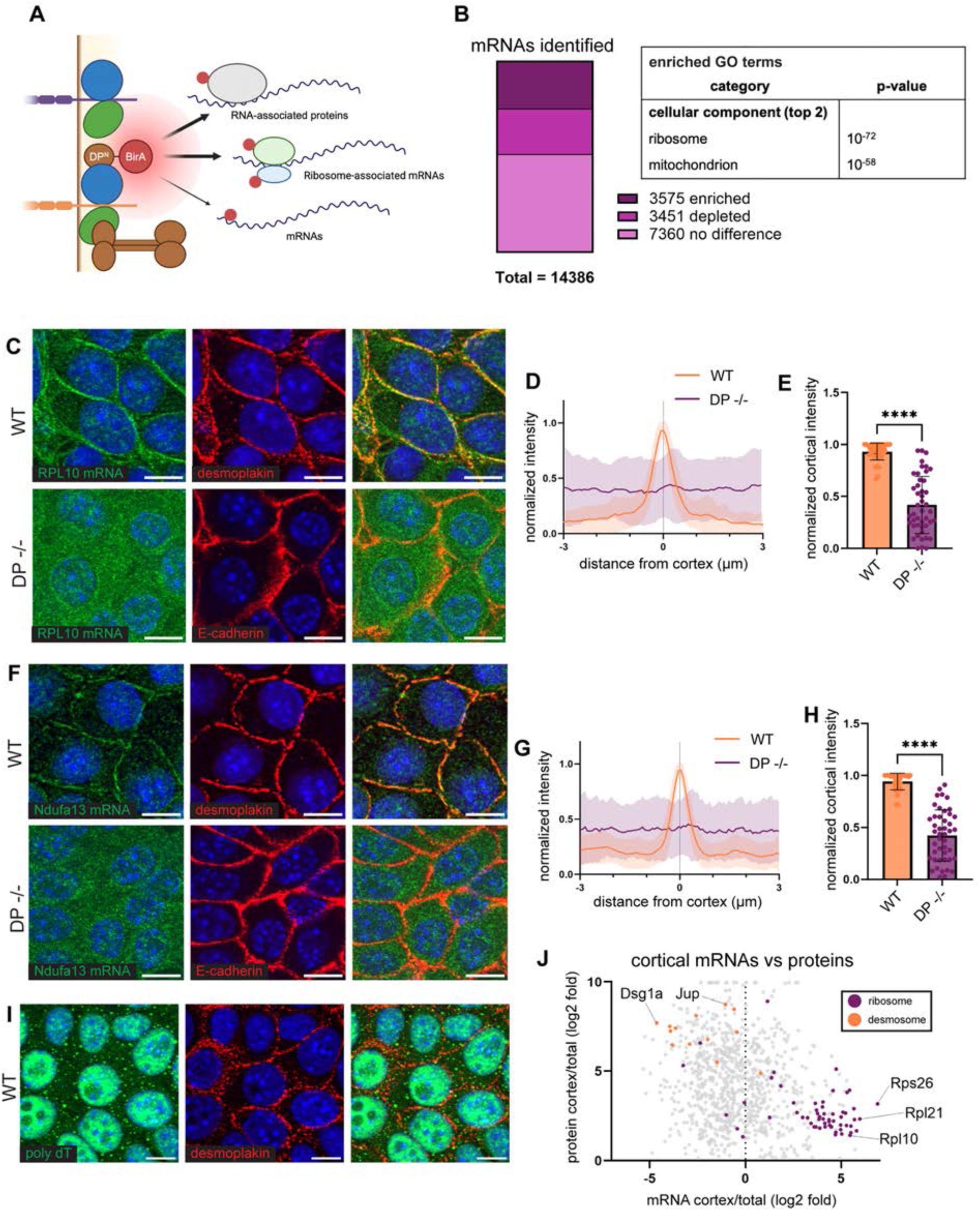
A subset of mRNAs is enriched at the cell cortex. (A) Schematic of BioID screen to identify mRNAs enriched at desmosomes. (B) Numbers of mRNAs identified as enriched or depleted from desmosomes and GO terms (cellular component) associated with enriched mRNAs. (C) smFISH labeling of RPL10 mRNA and desmoplakin or E-cadherin immunofluorescence in wild-type and desmoplakin-null mouse keratinocytes (scale bar = 10 µm). (D) Line scans of RPL10 fluorescence intensity measured across the cell cortex. Shaded area represents the standard deviation (WT n = 45 cells, DP-/- n = 45 cells; 3 biological replicates). (E) Relative intensity of RPL10 fluorescence at the center of the cell-cell boundary (WT n = 45 cells, DP-/- n = 45 cells; 3 biological replicates; p < 0.0001). (F) smFISH labeling of Ndufa13 mRNA and desmoplakin or E-cadherin immunofluorescence in wild-type and desmoplakin-null mouse keratinocytes (scale bar = 10 µm). (G) Line scans of Ndufa13 fluorescence intensity measured across the cell cortex. Shaded area represents the standard deviation (WT n = 45 cells, DP-/- n = 45 cells; 3 biological replicates). (H) Relative intensity of Ndufa13 fluorescence at the center of the cell-cell boundary (WT n = 45 cells, DP-/- n = 45 cells; 3 biological replicates; p < 0.0001). (I) Poly-dT FISH and desmoplakin immunofluorescence in wild-type mouse keratinocytes. (J) mRNA and protein enrichment at the cell cortex (PCC = -0.22, n = 925, p < 0.0001).

The cortical pool was highly enriched for mRNAs that encode for ribosomal and mitochondrial proteins (those that are nuclear encoded) (Fig. 4B). The pool of depleted mRNAs, in contrast, was enriched for transcripts encoding nuclear and ER proteins. Notably, transcripts encoding desmosomal proteins tended to be depleted from the cell cortex (Supplemental Table 1).

We turned to single molecule fluorescence in situ hybridization (smFISH) to validate the localization of mRNAs identified as being enriched at the cell cortex. We chose to probe for RPL10 and Ndufa13 mRNAs since they are abundant, long enough for probe tiling, and represent ribosomal and mitochondrial mRNAs, respectively, which were highly represented in our RNAseq dataset. We found that both mRNAs were enriched at the cortex (Fig. 4C-H). In contrast, probing for GAPDH or Ipo7 (cortically depleted), revealed no cortical enrichment, indicating that only select mRNAs are cortically enriched (Fig. S5B,C).

Given the requirement for desmosomes in ribosome localization, we next asked whether they were also required for mRNA localization. In desmoplakin null cells neither RPL10 nor Ndufa13 mRNA were cortically localized (Fig. 4C-H). While desmoplakin was required for mRNA cortical localization, keratins were dispensable, as mRNAs remained at the cortex in cells lacking keratin filaments (Fig. S5D). Together, our data demonstrate that the roles of desmosomes in localizing mRNA and regulating translation are independent of their role in anchoring intermediate filaments.

To further address molecular requirements for mRNA localization, we transfected desmoplakin null cells with various functional domains of desmoplakin to rescue mRNA localization. All constructs which include the N-terminal residues 1-584 (the minimal unit that targets to the cell cortex) were able to rescue mRNA localization (Fig. S6). This region is known to bind other desmosome components, but without keratin binding, desmosomes remain small and punctate in cells expressing this construct ^26,35–37^ . Therefore, the domains of desmoplakin that recruit ribosomes and Ndufa13 mRNA to the cell cortex are distinct. Further work will be needed to determine whether other cortically enriched mRNAs show a similar requirement.

To further address the molecular mechanisms for cortical mRNA localization, we first examined the role of cytoskeletal networks. Both actin and microtubules are known to be involved in actively transporting and/or anchoring mRNAs ^38–40^. Treatment of cells with nocodazole, cytochalasin B, Taxol, or blebbistatin for one hour did not disrupt the localization of Ndufa13 mRNA (Fig. S5D), demonstrating that the maintenance of this localization is not dependent upon the cytoskeleton. In addition, the localization of this mRNA did not depend upon its translation, as translational inhibitors (both puromycin and CHX) did not affect cortical enrichment (Fig. S5D). Whether these molecular requirements are similar for all cortically enriched transcripts will require further investigation.

In neuronal axons, mRNA stability correlates with localization ^11,41^. We thus analyzed our dataset to determine whether there were mRNA characteristics that were associated with cortical enrichment. Upon comparing our dataset to a published dataset of mRNA stability in embryonic stem cells, we found that cortically enriched mRNAs tended toward greater stability (Fig. S7A) ^42^. In addition, there is a known correlation between mRNA length and stability, with shorter transcripts being more stable ^43–45^ . Notably, we found a strong correlation of mRNA length to cortical localization, with cortically enriched transcripts being shorter compared to non-enriched and depleted mRNAs (Fig. S7B). Even within the enriched pool, there was a correlation between level of enrichment and mRNA length (Fig. S7E). In addition to total mRNA length, the length of the coding region and UTRs were also highly correlated with cortical enrichment (Fig. S7C,D). However, the molecular mechanisms that may mediate length and/or stability-dependent cortical localization remain unknown.

mRNAs for mitochondrial and ribosomal proteins are enriched in the axons of neurons, prompting us to broadly compare mRNA enrichment between the keratinocyte cortex and the axon ^46^. While these two categories were enriched in both, the two localized transcriptomes shared few other similarities, indicating the mechanisms that regulate mRNA localization in different cell types may be unique (Fig. S7F-H). Similarly, mRNAs enriched at the apical/basal sides of enterocytes showed no significant overlap with cortically enriched mRNAs in keratinocytes (Fig S7I-K) ^7^.

### Analysis of the cortical translatome reveals translational repression of many cortically-enriched transcripts

The predominantly proposed purpose for localizing transcripts is to bias the sites of their translation, allowing cells to produce proteins where they are needed ^47^. To globally assess this possibility, we compared the protein enrichment and the mRNA enrichment at the cortex (Fig. 4J). This analysis demonstrated that the localization of most transcripts does not correspond to protein localization. For example, while desmosomal proteins are highly enriched at the cortex, their mRNAs are depleted. The clear exception to this is ribosomes, whose mRNAs and proteins are both enriched at the cortex. However, whether this reflects local translation is unclear, especially as ribosomes are canonically assembled in the nucleolus, although some aspects of local assembly have been reported in other cell types ^48^. These data suggest that the cortical mRNA localization of many transcripts in keratinocytes serves a function other than protein localization.

Beyond coding functions, mRNAs can play structural roles in the cell, such as at focal adhesions and within the Golgi ^49 50^. Therefore, we treated cells with RNase A and examined desmoplakin localization. We verified loss of poly-dT signal upon RNase treatment, but did not observe any significant difference in desmoplakin localization or organization (Fig. S5A). These data demonstrate that mRNAs do not play an acute structural role in desmosome organization. Consistent with the strong enrichment of specific categories of mRNAs, this supports a non-structural role for cortical mRNAs and suggests a role for localization in translational regulation. That said, we cannot rule out more subtle structural roles for cortical mRNAs.

One possible reason for the lack of correlation between the cortical transcriptome and proteome is that the transcripts enriched at the cortex don’t reflect those being translated there. Thus, we sought to define the cortical translatome. To do this, we used DP-BirA expressing cells to biotinylate cortical proteins, including ribosomes (as diagramed in Fig. S4). Translating ribosomes were isolated from cell extracts using sucrose gradients, and the cortical, biotinylated molecules were purified. RNA-Seq revealed that the cortical and total pools of translation were distinct (Fig. 5A). These data demonstrate an unexpected compartmentalization of translation between the cortex and cytoplasm. There were approximately 2000 mRNAs whose presence on polysomes were enriched at the cortex compared to the total polysome pool while a similar number of mRNAs were depleted from the cortex (Supplemental Table 1).

**Figure 5:**
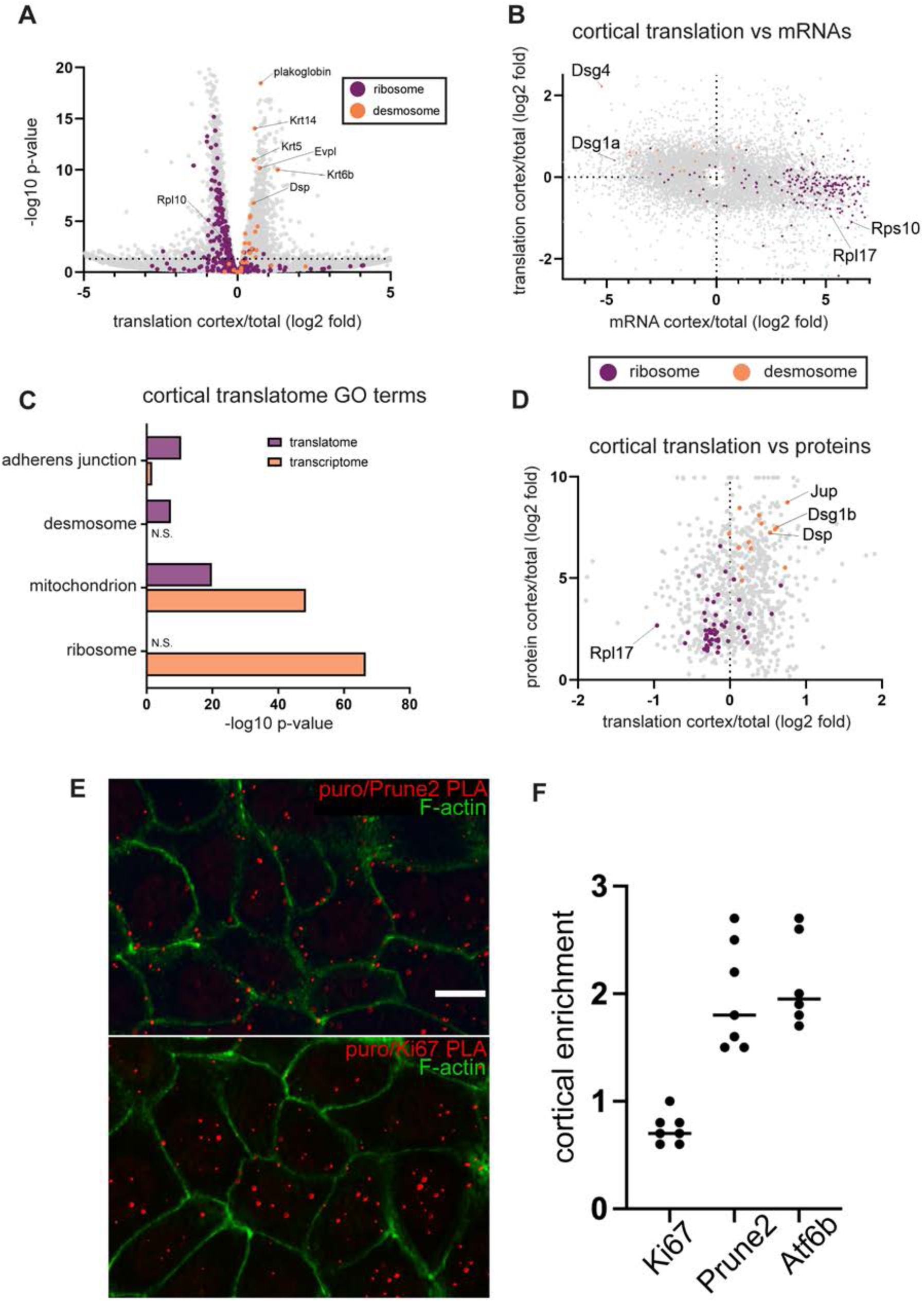
The cortical translatome is distinct from the cortical transcriptome. (A) Cortical translatome represented as a volcano plot with groups of desmosomal (orange) and ribosomal (purple) genes highlighted. (B) mRNA localization and translation at the cell cortex (PCC = -0.17, n = 7396, p < 0.0001). (C) Top cellular component GO terms within the cortical transcriptome and translatome. (D) Translation and protein localization at the cell cortex (PCC = 0.12, n = 877, p = 0.0005). (E) Images of a puro-PLA reaction for Prune2 and Ki67 showing puncta reflecting active translation in red. The cell cortex is marked by phalloidin staining of F-actin (green). (F) Quantitation of the cortical enrichment of translation for Ki67, Prune2, and Atf6b.

Overall, there was a significant, negative correlation between cortical translation and cortical localization of mRNAs, indicating that about 89% of cortical mRNAs are translationally repressed (Fig. 5B). While the list of cortically localized mRNAs was dominated by ribosomal proteins and mitochondria, the list of cortically translating mRNAs was highly enriched in adhesion and cytoskeleton proteins (Fig. 5C). None of the ribosomal mRNAs were enriched in translation at the cortex, while subsets of mitochondrial mRNAs were either depleted from the cortex (n=379, p-value 10^−101^) or enriched there (n=221, p-value 10^−20^). Thus, while mitochondria and ribosomes dominated the cortical transcriptome, cell adhesion structures, including all core components of the desmosome, were enriched in the cortical translatome (Fig. 5B,C). This list also included many keratins (Krt5/14/6a/6b/15/16/17/78/80), perhaps consistent with the idea that keratin assembly occurs near the cortex ^51^. Together, our data demonstrate that there is active translation at the cortex of many non-cortically enriched transcripts (along with about 11% of the cortically enriched transcripts), while a large fraction of the cortically-enriched transcripts are translationally repressed at this site.

Given these results, we compared the cortical translatome with protein localization and found a much stronger correlation between these two than between the cortical proteome and the cortical transcriptome (compare Fig. 5D to 4J). These data highlight the importance of local translational regulation and demonstrated the utility of examining local translatomes as opposed to transcriptomes. That said, there are still significant differences between the cortical translatome and the cortical proteome. Overall, our data suggest that while there is active translation occurring at the cortex, there is also transcript-specific regulation as the most enriched transcripts are translationally repressed.

We validated the translatome results using proximity ligation assays with puromycin (puro-PLA). We chose two candidates, Prune2 and Atf6b, based on their high cortical enrichment, available antibodies that recognized their amino-terminal region, and their large size (longer residence time on the ribosome). For each of these nascent proteins, we found enrichment at the cell cortex, while puro-PLA for an mRNA that was not cortically enriched, Ki67, was more uniformly distributed (Fig. 5E,F). The cortical signal is not due to preferential localization of either of these proteins to the cortex as Prune2 is keratin-associated while Atf6b is nuclear-enriched.

### Cortical localization of RISC is desmoplakin-dependent

We next wanted to further explore mechanisms of translational repression at the cortex. As there is a diverse pool of mRNAs localized here as well as many RNA binding proteins (RBPs) and translational regulators, there are likely multiple mechanisms of regulation. One intriguing candidate was the RNA-induced silencing complex (RISC), which has known roles in translational repression, and which was identified in our proteomic analysis (Fig. S8A,B). RISC functions to either degrade or translationally repress mRNAs ^52^. Phosphorylation of Argonaute 2 (Ago2) at S387 shifts RISC mRNA regulation towards repression ^53^. We saw clear enrichment of both Ago2 and pAgo2^S387^ at the cortex, suggesting that at least some of the cortical RISC was involved in translational repression (Fig. 6A-C; S8C,D). Cortical localization of Ago2 and pAgo2^S387^ was dependent upon desmoplakin, but not the adherens junction protein p120-catenin (Fig. 6A-E;S8C,D). Ago2 localizes to zonula adherens in simple epithelia in a p120-catenin-dependent manner ^54^, thus these data demonstrate a distinct mechanism of cortical recruitment in keratinocytes. Cortical Ago2 and pAgo2^S387^ localization was also observed in intact tissue, where it was again dependent upon desmoplakin (Fig. 6F,G, and Fig. S8E,F).

**Figure 6:**
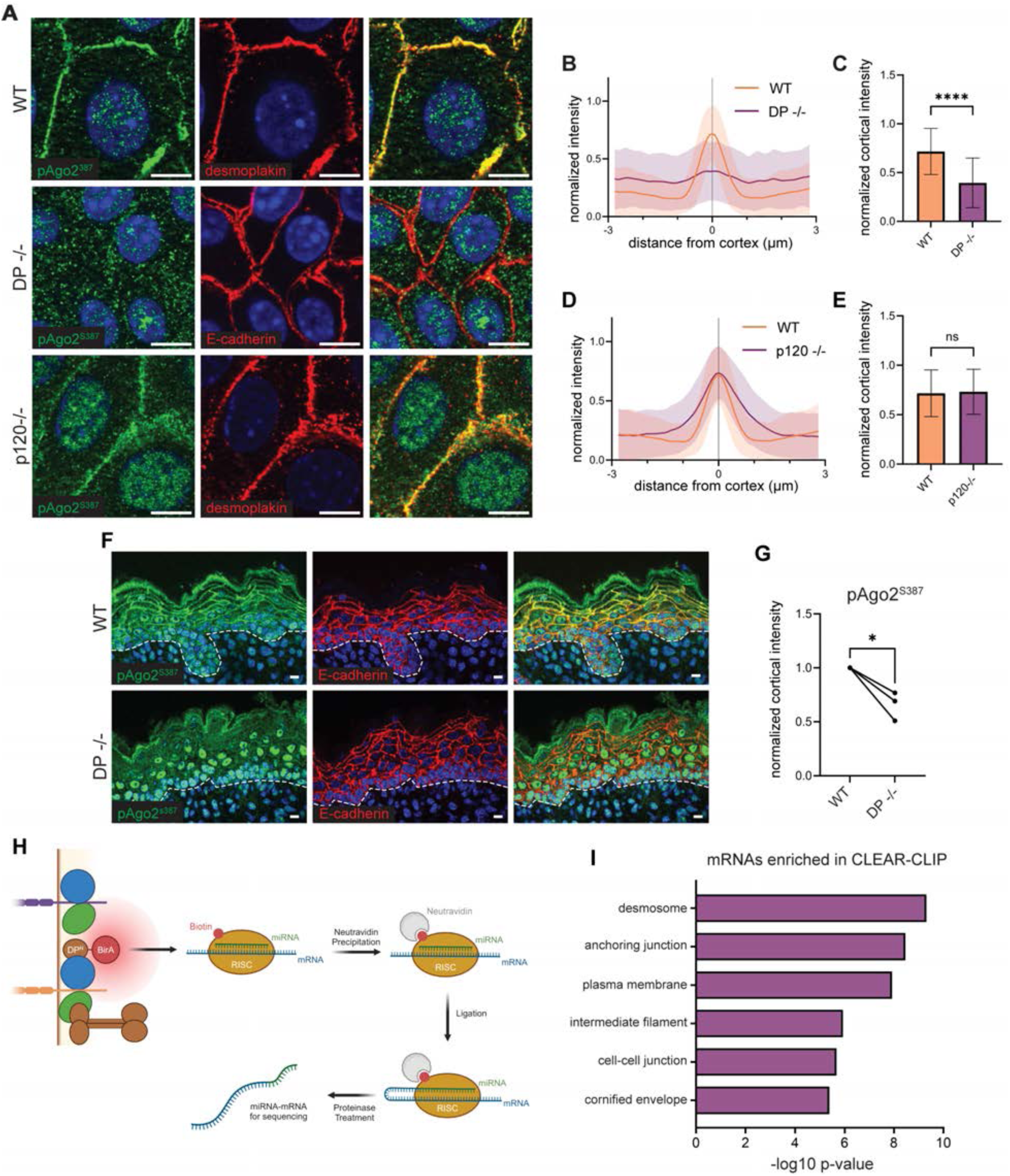
Ago2 localizes to the cortex and associates with select mRNAs. (A) Representative immunofluorescence images of pAgo2^S387^ in wild-type, desmoplakin-null, and p120-null mouse keratinocytes counterstained with desmoplakin and E-cadherin (scale bar = 10 μm). (B, D) Line scan analysis of pAgo2^S387^ over cell-cell boundaries comparing (B) wild-type and desmoplakin-null and (D) wild-type and p120-null. Shaded region represents the standard deviation from the mean (WT n = 558 cells; DP-/- n = 528 cells; p120-/- n = 449 cells; 3 biological replicates). (C, E) Relative intensity of pAgo2^S387^ at the center of the cell-cell boundary comparing (C) wild-type and desmoplakin-null (p < 0.0001) and (E) wild-type and p120-null (p = 0.3029). (F) Representative immunofluorescence images of pAgo2^S387^ and E-cadherin in back skin from E18.5 wild- type and desmoplakin-null mouse embryos; basement membrane indicated by a white dotted line; scale bar, 10 μm. (G) Relative intensity of pAgo2^S387^ at the center of the cell- cell boundary from tissue comparing wild-type and desmoplakin-null (p = 0.0466, n = 3 mice). (H) Schematic of CLEAR-CLIP technique for identifying mRNAs associated with RISC at the cell cortex. (I) Select cellular component GO terms for the top 200 mRNAs identified by CLEAR-CLIP as being associated with RISC at the cell cortex.

To better understand cortical RISC function, we set out to identify the mRNAs and miRNAs associated with the cortical pool. Using Dsp-BirA expressing cells, we biotinylated cortical proteins, including RISC, isolated over streptavidin, and performed CLEAR-CLIP analysis which allows identification of miRNA-mRNA pairs that have been ligated together ^55^ (Fig. 6H). Remarkably, we found that cell adhesion, cytoskeleton, and epidermal barrier (cornified envelope) genes were among the highest represented in this group (Fig. 6I). Lists of the most abundant mRNAs and miRNAs can be found in Supplemental Table 1.

### Wounding induces RISC delocalization and translational responses

The types of mRNAs associated with cortical RISC led us to hypothesize that these complexes serve as a reservoir for mRNA transcripts to promote/restore epithelial integrity when needed. In support of this possibility, we found that there was a loss of cortical Ago2 in response to wounding of cultured cells (Fig. 7A,B). This was also observed in wounded skin, although in this case we stained for pAgo2^S387^ as this antibody works better in tissue (Fig. 7D,E). Further, perturbing desmosomes with an antibody that binds to Dsg3 (monoclonal AK23 antibody, which mimics some of the effects of antibodies found in the autoimmune disease pemphigus vulgaris ^56^) also resulted in a decrease in cortical pools of Ago2 (Fig. 7G,H). In the cases of *in vitro* wound healing and pemphigus antibody treatment, loss of Ago2 was not associated with loss of cortical DP, suggesting that the interaction is regulated and not constitutive.

**Figure 7:**
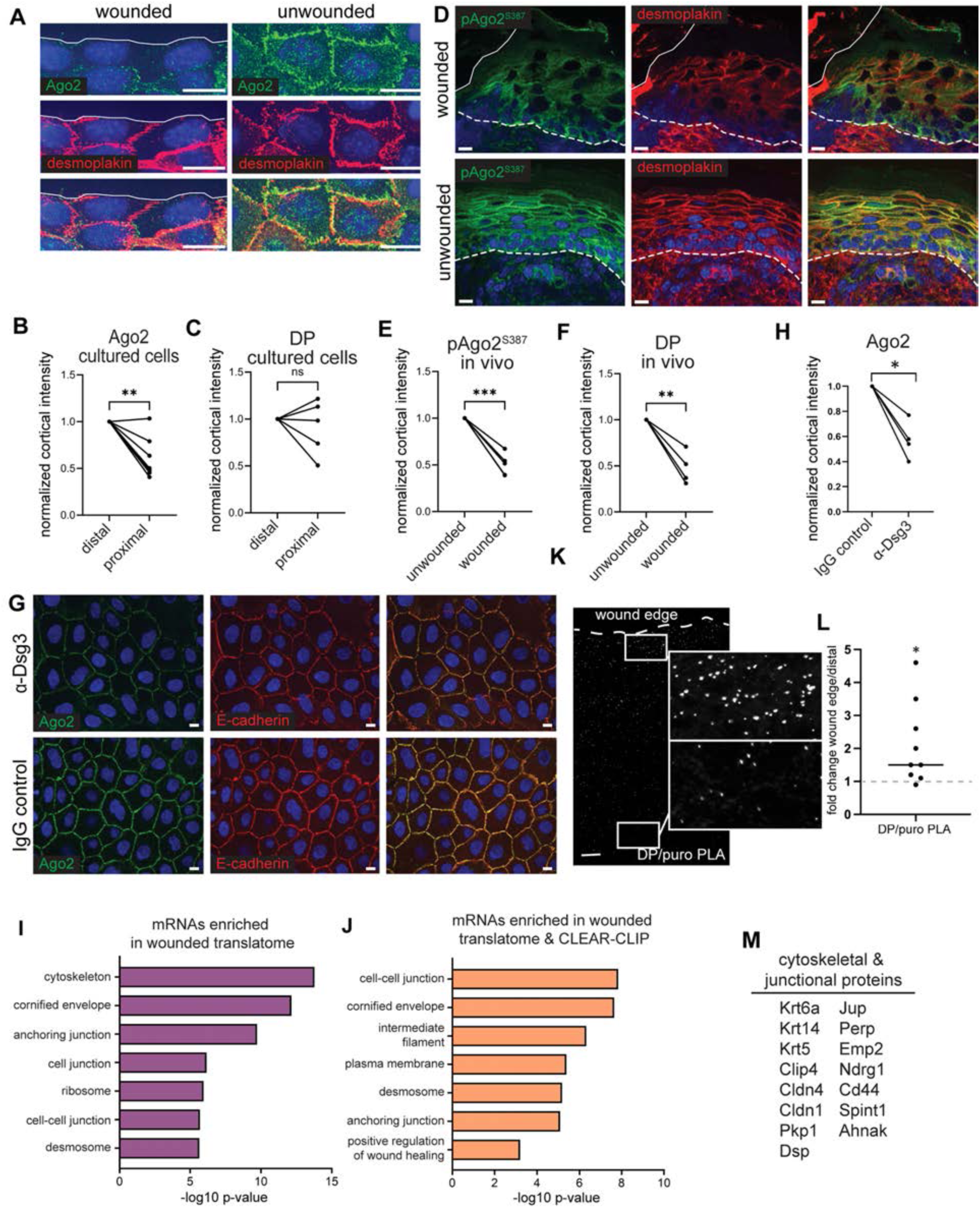
Delocalization of Ago2 following desmosome perturbation. (A) Confluent monolayers of mouse keratinocytes were scratched with a pipetted tip and allowed to recover from injury for 1 hour, stained for Ago2 and desmoplakin. Wounded edge (solid line) and the unwounded section taken 2 fields of view away (scale bar = 10 μm). (B, C) Relative intensity of (B) Ago2 and (C) desmoplakin at the center of the cell-cell boundary from *in vitro* scratch experiment from (A) (Ago2 n = 7 biological replicates, p = 0.0041; DP n = 5 biological replicates, p = 0.5505). (D) Representative immunofluorescence images of pAgo2^S387^ in back skin collected from P0 mouse pups that were wounded with a 0.5 cm biopsy punch and allowed to recover for 1 hour. Solid line marks the edge of the wound. Basement membrane indicated by a white dotted line (scale bar = 10 μm). (E, F) Relative intensity of (E) pAgo2^S387^ and (F) desmoplakin at the center of the cell-cell boundary from *in vivo* wound experiment from (D) (pAgo2^S387^ n = 5 mice, p = 0.0007; DP n = 4 mice, p = 0.0098). (G) Representative immunofluorescence images of Ago2 and E-cadherin in primary human keratinocytes that were either treated with anti-Dsg3 (AK23, pemphigus mimic) antibodies or a control IgG (scale bar = 10 μm). (H) Relative intensity of Ago2 at the center of the cell-cell boundary in pemphigus antibody exposure experiment from (G) (n = 4 biological replicates). (I) Select cellular component GO terms for mRNAs identified as being upregulated in the translatome following wounding. (J) Select cellular component and biological process GO terms for mRNAs which were both among the top 200 mRNAs associated with RISC at the cell cortex and upregulated in the translatome following wounding. (K) Proximity ligation assay for desmoplakin and puromycin 40 minutes after wounding. (L) Quantitation of fold difference of puro-PLA puncta at the wound edge verses sites distal to the wound (n = 9 images from 3 biological replicates, p=0.0176). (M) Select genes from the overlap of mRNA upregulated in the translatome following wounding and among the top 200 associated with RISC at the cell cortex.

Due to these findings, we repeated our analysis of the cortical translatome, but this time we scratch-wounded the cells to determine if the translatome was altered. Notably, we found that wounding induced a reproducible increase in disomes, as well as a smaller increase in 3+ ribosomes, suggesting a wound-induced translational response (Fig. S9A-C). GO analysis of genes that were translationally upregulated revealed roles in cytoskeleton, cornified envelope formation, and adhesion – processes important for re-establishing epithelial integrity and barrier formation (Fig. 7I). Importantly, there were few genes in this list that were also changed at the transcript level ^57^, and these were removed from the analysis.

We next compared the cortical/cytoplasmic ratio of translating transcripts in control and wounded epithelial cell layers to determine whether changes were related to spatial location. We found that transcripts that were either most highly translationally active at the cortex and those that were most translationally active in the cytoplasm were the ones most likely to be changed (Fig. S9D). While the underlying mechanism is unclear, this demonstrates significant translational changes upon wounding.

We next compared the mRNAs whose translation was upregulated upon wounding with the mRNAs associated with cortical RISC. There was significant overlap between these lists, and they were again enriched for adhesion, cytoskeleton and barrier formation genes (Fig. 7J,K). These included many desmosomal and tight junction components, notably desmoplakin itself. We therefore performed puro-PLA on desmoplakin under wounding conditions and noted an upregulation of translation at the wound site as compared to more distal regions (Fig. 7K,L). These data are consistent with the idea that at least some of the mRNAs sequestered by cortical RISC are released for translation upon wounding.

## DISCUSSION

Our data reveal that translation is compartmentalized between the cortex and the cytoplasm in epidermal cells. This occurs at multiple levels -ribosomes, mRNAs, and RNA-binding proteins like RISC.

Ribosomes localize to distinct subcellular structures in a number of cell types ^7,8,58^, however, the molecular mechanism of their localization is largely unknown. In the case of cardiomyocytes, ribosomal protein SA (RPSA) is required for ribosome localization to the sarcomere, however additional factors are unknown ^59^. We demonstrate that the core desmosomal protein, desmoplakin, is required for cortical recruitment of ribosomes in keratinocytes. Further, we identified a minimal domain, PRD-A, that was necessary and sufficient for ribosome recruitment. PRDs are intermediate-filament binding domains; however, PRD-A has weak affinity for keratins and when expressed in cells, does not decorate filaments ^29,37,60^. It also has a notably more basic characteristic than other PRDs, which could be important for interaction with the acidic rRNA of ribosomes ^61^. Desmoplakin is not only found at the cortex in keratinocytes, but also the intercalated discs of cardiomyocytes, where ribosomes also accumulate, suggesting this function may be conserved in other tissues ^8^. Further, the protein dystonin also has a very similar PRD-A domain. Dystonin localizes to the Z-band in striated muscle, another site of ribosomal localization, and thus PRD-A domains have the potential to mediate ribosome localization in diverse tissues ^62^. Notably, keratins and desmosomes serve as attachment sites for endoplasmic reticulum (ER) at the cell cortex ^16^. While rough ER could serve as part of the mechanism for localization of ribosomes to the cortex, our finding that keratins and microtubules are not required for ribosome localization or cortical translation suggests that at least some of this occurs independently of the ER (as ER organization is dependent on both cytoskeletal networks) ^14^.

There was also an unexpected and broad compartmentalization of mRNAs in epidermal cells. While mRNA localization in polarized cells is well-studied, this cortical compartmentalization has not been fully appreciated before. In the differentiated epidermis where this localization occurs, cells are not polarized. Only a few transcripts, including dlg1 in C. elegans, are known to be localized to the cortex where they are actively translated ^63^. In epidermal cells, we found that half of transcripts were either enriched or depleted from the cortex, thus, this appears to be a major axis of organization in these cells. This function is dependent upon desmoplakin but is molecularly separable from ribosome recruitment. While prior work has demonstrated some interactions between desmosomal components and mRNA/translational machinery, this was thought to occur outside of the desmosome and to be independent of it ^64,65^. Our proteomic analysis of desmosomes revealed a significant number of RNA-binding proteins (RBPs) ^21^, providing candidates that may recruit distinct classes of mRNAs. It is therefore likely that there are multiple factors that participate in cortical mRNA localization and regulation. Notably, there was a striking enrichment of short mRNAs at the cell cortex. This pattern was also observed in axons, where the correlation between small size and increased stability of mRNAs was proposed to mediate the localization, as only stable transcripts had sufficient time to diffuse into axons ^11,41^. The small size of keratinocytes makes this explanation unlikely in the skin, suggesting novel mechanisms linking mRNA length to localization.

Our generation of datasets for the proteome, the transcriptome, and the translatome at the cell cortex allowed us to directly address roles for transcript localization and translation. Surprisingly, we found that most transcripts localized at the cell cortex were not actively translated at this site under homeostatic conditions. In contrast, there was a much better correlation between cortical translation and cortical protein localization, although there were many examples of mRNAs/proteins that do not follow this pattern. This work highlights two important points: there is risk in assuming that the function of mRNA localization is solely for local translation, and that there are layers of local translational regulation that determine which mRNAs are translated and under what conditions. Further the use of local biotinylation with polysome analysis is expected to be a broadly useful technique to determine local translatomes.

The presence of translationally repressed transcripts at the cortex suggests the presence of mRNA binding proteins that control translation. Indeed, our BioID proteomics revealed many RBPs and translational regulators enriched at the cortex, including RISC. RISC localization was also dependent upon desmoplakin, but distinct from ribosomes and mRNAs, its cortical localization was decreased by both wounding and pemphigus antibodies. Analysis of mRNAs bound to cortical RISC revealed transcripts enriched for cell adhesion and wound healing genes. Notably, when we examined translational changes in response to wounding, many of these cell adhesion genes were translationally upregulated. These data suggest that cortical RISC (and potentially other RBPs) may serve as a reservoir for mRNAs that can be rapidly mobilized for translation in response to integrity defects or other needs. Further, rather than acting just as passive adhesions, this work raises the possibility that desmosomes are active sensors of epidermal integrity that activate translation responses in response to injury.

Limitations of the study: While we demonstrate that transcripts associated with RISC are upregulated upon wounding, we are not able to demonstrate that the specific mRNAs bound by RISC are released and then translated. In addition, while we identify translational repression of many transcripts at the cell cortex, the mechanism underlying this repression (beyond RISC-bound transcripts) remains unknown; nor are the stimuli that result in activation of translation of these repressed transcripts. While we identify a role for desmoplakin in recruitment of specific mRNAs to the cell cortex, further work is needed to establish the RBP intermediaries required for this recruitment. Finally, desmosomes are not uniform throughout epidermal differentiation and we do not address how the local transcriptome and translatome may change through differentiation.

## METHODS

### Cell culture

All cell culture studies were performed on mouse or human keratinocytes. All keratinocyte cell lines were grown at 37°C and 7.5% CO_2_. Mouse keratinocytes were isolated from the backskin of E18.5 embryos by dispase treatment and trypsinization. After several passages on mitomycin-treated fibroblast feeders, keratinocyte lines were grown in the absence of feeders in E no Ca^2+^ media (3:1 DMEM:F12 [Invitrogen], 0.5 µg/mL insulin [Sigma, 12585-014], 0.1 nM cholera toxin [MP, 02150005], 0.5 µg/mL transferrin [Sigma,], 0.4 µg/mL hydrocortisone [Calbiochem, 386698,], 0.2 µM T3 [Sigma T2752, 0.2], 15% fetal bovine serum [Hyclone, A-1115-L]), which was first chelated with Chelex (Bio-Rad, 142-2842) to remove calcium and afterward supplemented to a final concentration of 0.05 mM with CaCl_2_). To establish stable DP(1-584)-BirA cell lines, mouse keratinocytes were infected with lentivirus, followed by selection in puromycin (2 µg/mL). Unless otherwise indicated, mouse keratinocytes were cultured in E no Ca^2+^ media until confluency, at which point CaCl_2_ (1.2 mM) was added to induce differentiation and cell-cell adhesion. HaCaT cells were cultured in DMEM + 10% unchelated FBS (Hyclone, A-1115-L). Human primary cells were gifts from the Jennifer Zhang lab (Duke University Medical Center, Department of Dermatology) and were cultured in Keratinocyte SFM (Thermo, 17005042) until confluent, at which point CaCl_2_ (1.2 mM) was added to induce differentiation and cell-cell adhesion. Primary cells for pemphigus experiments were treated with mouse anti-Dsg3 (MBL, AK23, D219-3) for 1 hr at 10 µg/mL. DNA transfections were performed using TransIT-LT1 (Mirus, MIR 2304), polyethyleneimine, or Cellfectin II (Thermo, 10362100)

### Lentivirus generation and infection

Lentiviruses were made by co-transfection of pMD2.G (Addgene, 12259), psPAX2 (Addgene, 12260), and pLIX402-DP(1-584)-BirA-HA into HEK-293T cells using polyethyleneimine. Supernatant was collected at 24‒96 hrs after transfection. Lentivirus was concentrated with PEG-8000 buffer (40% PEG-8000, 1.2 M NaCl), resuspended in 1X PBS, flash-frozen, and stored at -80°C until use. On the day of infection, 1 aliquot of lentivirus was added to mouse keratinocytes grown to 50-60% confluency in a 3.5 cm dish. Polybrene (6 µg/mL) was added to the culture media. The infection mixture was incubated with the cells for 8 hrs, and then the media was replaced. Cells were split upon reaching confluency and then 48 hrs after infection were treated with puromycin (2 µg/mL). For several weeks, cells were passaged in puromycin to select for stably expressing cells.

### Microscopy

Cell stains and tissue sections were imaged on a Zeiss AxioImager Z1 microscope with an Apotome 2 attachment using a Plan-Apochromat 63x/1.4 oil DIC objective, Axiocam 506 mono CCD camera, and Zen Blue 2.60 software (Zeiss). Super-resolution images were taken on a Zeiss Elyra 7 system with a Plan-Apochromat 63x/1.4 oil DIC M27 objective, dual pco.edge 4.2 CL HS sCMOS cameras, and Zen Black 3.10 software.

### Immunofluorescence

Mouse keratinocytes were grown to confluency on glass coverslips in E no Ca^2+^. Once confluent, CaCl_2_ (1.2 mM) was added to initiate differentiation and adhesion. Cells were grown in calcium for 24 hrs, and then were briefly washed in 1X PBST (0.2% Triton X-100) to pre-permeabilize, followed by fixation in cold MeOH at –20°C for 3 min.

Coverslips were washed twice in PBST and then blocked with buffer (3% BSA, 5% NGS, 5% NDS) for 15 min. Primary antibodies were diluted in blocking buffer and added to the coverslips to incubate for 1 hr at room temp or overnight at 4°C. Next, the coverslips were washed 3 times in PBST, followed by incubation with secondary antibodies and Hoechst 33258 diluted in blocking buffer for 30 min. The coverslips were then washed 3 more times and mounted to slides using ProLong Gold (Thermo, P36930) or antifade. For Ago2/pAgo2^S387^ tissue stainings, backskin was collected from P0 pups and fixed in 4% PFA for 30 min, then mounted in OCT (Sakura 483).

Cryosections were taken at 10 µm and adhered to Superfrost Plus slides (VWR, 48311-703), followed by drying for 2 hrs at room temp and a post fix in 4% PFA for 15 min. For RPL10A tissue stainings, whole E18.5 embryos were mounted unfixed in OCT and cryosections were taken at 10 µm and adhered to Superfrost Plus slides followed by drying for 2 hrs at room temp and a post fix in 100% MeOH for 15 min. Slides were then washed and permeabilized in PBST for 15 min, then blocked for 1 hr in blocking buffer. Antibody staining was performed as with coverslips. Primary antibodies used include: RPL10A (1:100, Abclonal A5925), RPS3 (1:100, Novus NBP1-33691), anti-PABP (1:100, Santa Cruz sc-32318), Desmoplakin (1:200, Millipore, MA1-83118), E-Cadherin (1:200, Invitrogen, 13-1900), GFP (1:1000, Abcam, ab13970), V5 (1:100, Cell Signaling 13202S), Ago2 (1:100, ThermoFisher, PA5-66129), pAgo2^S387^ (1:100, PhosphoSolutions, AP5291). Secondary antibodies used include: Donkey Alexa Fluor 488– and 647–conjugated series (Life Technologies/Thermo Fisher) and Donkey Rhodamine Red–conjugated series (Jackson Immuno-Research). All cell stainings were performed at minimum three times. Line-scans were performed in Fiji by drawing across cell-cell borders marked by a cortical marker (i.e. desmoplakin or E-cadherin). Data from each line was normalized to their minimum and maximum intensity. Graphs represent averages across 40-50 lines from 3 biological replicates. For co-localization/correlation in GEM imaging, regions of the cytoplasm (∼10 µm^2^) were sampled and Pearson’s correlation coefficient and Mander’s overlap coefficient were calculated in Zen Black 3.10. One region was selected per cell with >10 cells used per biological replicate. Unpaired T-tests were used to calculated statistical significance.

### Nascent peptide labeling

Differentiated mouse keratinocytes or HaCaT cells were cultured as stated before. For O-propargyl-puromycin (OPP) labeling, cells were treated with OPP (50 µM, Bioconjugate technologies, 1407-5) diluted in DMSO by adding it to the culture media for 5 min. For L-homopropargylglycine (HPG) labeling, cells were washed with 1X PBS several times, then placed in DMEM lacking methionine (Gibco, 21013024) to deplete methionine reserves. After a 1 hr methionine depletion, HPG (5 µM, Click Chemistry Tools, 1067-25) diluted in DMSO was added to the culture media for either 3 or 5 min (as indicated). For all nascent peptide labeling experiments, cells were rapidly washed at the end of the labeling period with 1X PBST, then fixed in 100% MeOH at -20°C for 2 min. After fixation, cells were washed several times in 1X PBST followed by incubation with Click reaction buffer (10 mM TBS pH7.4, 200 µM CuSO_4_, 2 μM MB488 or Alexa Fluor 647 azide, 400 μM TCEP) for 30 min at room-temp. Following Click-labeling, cells were washed several times in 1X PBST to remove excess dye, then immunostained as stated before.

### RNAseq

Stable DP(1-584)-BirA keratinocytes were cultured as above. Doxycycline (2 µg/mL) and CaCl_2_ (1.2 mM) were added to confluent cells to induce differentiation and expression of the DP(1-584)-BirA construct. At 24 hrs post doxycycline induction, biotin (100 µM) was added to cells which continued in culture for 24 hrs. After the biotin labeling period, cells were lysed in RIPA Buffer (50 mM Tris, pH 8, 1% Triton, 150 mM NaCl, 0.5% SDS, 50 mM Triton, 1 mM EDTA, Protease Inhibitor Cocktail [Roche, 11697498001], RNasin Plus [Promega N2611]), and incubated with pre-washed Neutravidin resin (Pierce/Thermo, 29200) overnight at 4°C. The following morning, the resin was washed three times in lysis buffer, and RNA was isolated from the resin using Qiagen Buffer RLT and RNase-free EtOH (E7148) and RNA was isolated according to the Qiagen RNeasy Mini kit (74104). Whole cell RNA was isolated from wild-type mouse keratinocytes incubated in Ca^2+^ for 24 hrs according to the Qiagen RNeasy Mini kit.

Biological triplicate samples were sent to Novogene (San Diego, California) where they were library prepped by poly-A purification, reverse transcription, and low-input amplification. Samples were sequenced in a single lane on a NovaSeq X Plus with a minimum of 20 million paired reads (150 bp) per sample. Following sequencing, data underwent Novogene’s data QC and was mapped to the mouse genome.

### Transcriptome analysis

Transcriptome data from Moor et al., 2017 and Zappulo et al., 2017 were compared to our transcriptome dataset. Genes were filtered for those represented in both lists. For the purposes of calculating Pearson’s correlation coefficients and statistical significance, only genes that were significantly enriched in either dataset were considered (p<0.05). Ribosome and mitochondrion genes were selected from the list based on cellular component GO term (https://david.ncifcrf.gov/home.jsp). Analysis was conducted in GraphPad Prism.

### smFISH probes

Oligo probes were designed using the Stellaris Probe Design tool (Biosearch Technologies). Each probe set was comprised of 24-48 probes of 20-nt each and was mapped to mouse cDNA sequences obtained through ENSEMBL (NIH). Probes were conjugated to Quasar570 or Quasar670 dyes. Each probe set was diluted to 12.5 µM and stored at -80°C until use.

### smFISH

Mouse keratinocytes were grown to confluency on glass coverslips in E no Ca^2+^ media. Once confluent, CaCl_2_ (1.2 mM) was added to initiate differentiation and adhesion. Cells were grown in calcium for 24 hrs, and then were briefly washed in 1X PBST (0.2% Triton X-100) to pre-permeabilize, followed by fixation in 4% paraformaldehyde for 10 min. Following fixation, coverslips were washed in 1X PBS, followed by an additional fixation and permeabilization in MeOH at -20°C overnight. Prior to hybridization, coverslips were washed with wash buffer (2X SSC, 10% de-ionized formamide) twice for 5 min. Oligo probes were diluted to 1.25 µM in hybridization buffer (2X SSC, 10% de-ionized formamide, 10% dextran sulfate). This mixture was applied to coverslips, which were transferred to a humidity chamber, and incubated at 37°C overnight. The following morning, coverslips were washed in wash buffer twice at 37°C for 30 min. Immunofluorescence was performed on the hybridized coverslips as stated above. All cell hybridizations were performed with a minimum of three biological replicates. Line-scans were performed in Fiji by drawing across cell-cell borders marked by a cortical marker (i.e. desmoplakin or E-cadherin). Data from each line was normalized to their minimum and maximum intensity. Graphs represent averages across 40-50 cell boundaries from 3 biological replicates. Using GraphPad Prism, an unpaired T-test was used to calculated statistical significance.

### Polysome profiling and isolation

For cortical ribosome isolation, DP(1-584)-BirA keratinocytes were grown to confluency in 10 cm dishes in E no Ca^2+^ media. Once confluent, doxycycline (2 µg/mL) and CaCl_2_ (1.2 mM) and was added to initiate differentiation and adhesion. Cells were grown in calcium for 24 hrs, followed by addition of cycloheximide (200 µM) for 5 min to stall ribosomes. Dishes were washed in cold 1X PBS (with 200 µM cycloheximide) followed by lysis in cold buffer (200 mM KCl, 25 mM K-HEPES pH 7.4, 15 mM MgCl_2_, 1% NP-40S, 0.5% sodium deoxycholate, 200 µM cycloheximide, 1 mM DTT, 40 U/mL RNasin Plus [Promega, N2611], 2 mM PMSF, 1X Halt Protease/Phosphotase Inhibitor [Thermo, 78440]). Lysate was passed 25-30 times through a syringe with a 23-gauge needle and incubated on ice for 10 min. Lysate was then clarified by centrifugation at 21,000 xg for 10 min at 4°C, and supernatant was loaded onto a 15-50% sucrose gradient ((200 mM KCl, 25 mM K-HEPES pH 7.4, 15 mM MgCl_2_, 200 µM cycloheximide, 1 mM DTT, 10 U/mL RNasin Plus [Promega N2611]) in a 13.2 mL ultracentrifuge tube (Beckman, 331372). Samples were centrifuged at 35,000 rpm for 3 hrs at 4°C in an SW41 rotor. Following centrifugation, density gradients were fractionated into 0.5 mL aliquots using an Isco tube piercing system and peristaltic pump (Isco, 67-9000-177) with UA-6 Detector with 254 nm filter. For monosome/polysome blots, fractions corresponding to monosome or polysome peaks were pooled and diluted 2-fold in buffer without sucrose. These diluted fractions were incubated with pre-washed Neutravidin resin (Pierce/Thermo, 29200) overnight at 4°C. The following morning the resin was washed three times in 50 volumes of lysis buffer for 5 min and then boiled in 2X SDS loading buffer to elute protein. Meanwhile, the depleted supernatant from the beads was precipitated and concentrated using trichloroacetic acid (TCA, Sigma, T6399) overnight at 4°C, washed several times in 70% EtOH, then resuspended, sonicated, and boiled in 2X SDS loading buffer. Samples were loaded onto a 10% SDS-PA gel to separate proteins, which were then transferred onto a nitrocellulose membrane for blotting. Blots were dried, then rehydrated and blocked with 3% milk (in PBS + 0.1% Tween-20) for 30 mins, incubated with primary antibody (diluted in 5% BSA in PBST) for 2 hrs at room-temp, washed 3 times with PBST, incubated with the secondary antibody for 1 hr (diluted in 5% BSA in PBST), then washed 3 times in PBST. Images were acquired on a LI-COR Fc system using a 700 nm laser line with 2 min integration times. Band intensity was quantified in Image Studio version 5.2 with background subtraction. Experiment was repeated in biological triplicate. An unpaired T-test was used to calculated statistical significance in GraphPad Prism. For PRD-A expression and polysome profiling, HEK293T cells in 10 cm plates were transfected with 5 µg of pEF6-V5/HIS-TOPO-DP(2004-2233) using PEI (3:1). Cells were cultured for 48 hrs after transfection, at which point CHX was added (as stated before), cells were washed with cold PBS (with CHX) and subsequently lysed in cold lysis buffer. For EDTA treatment, 30 mM EDTA was added to the lysate. For RNase A treatment, 1 mg/mL was added to the lysate, which was then incubated for 20 min. The lysates were clarified by centrifugation and loaded on to sucrose gradients (15-50%) and separated as before. From each gradient, 20 fractions were collected at 30 sec intervals (∼0.5 mL). Each fraction was precipitated with 20% TCA overnight at 4°C, washed in 70% EtOH several times, and sonicated and boiled in 2X SDS loading buffer. From each fraction, 10% of the total was loaded onto a 10% SDS-PA gel to separate proteins, which were then blotted as stated before. Antibodies used include: RPL10A (1:100, Abclonal A5925), RPS3 (1:100, Novus NBP1-33691), V5 (1:100, Cell Signaling 13202S), and goat anti-rabbit IgG Alexa Fluor 680 (1:10000, Invitrogen A-21109).

### Translatome isolation and sequencing

Stable DP(1-584)-BirA keratinocytes were cultured as above. Doxycycline (2 µg/mL) and CaCl_2_ (1.2 mM) were added to confluent cells to induce differentiation and expression of the DP(1-584)-BirA construct. At 24 hrs post doxycycline induction, biotin (100 µM) was added to cells which continued in culture for 16 hrs. After the biotin labeling period, cells were lysed in polysome lysis buffer and lysates were clarified and run as stated above. For each sample, fractions containing monosomes and polysomes were pooled and diluted in 1 volume of buffer (200 mM KCl, 25 mM K-HEPES pH 7.4, 15 mM MgCl_2_, 200 µM cycloheximide, 1 mM DTT, 10 u/mL RNasin Plus [Promega, N2611]) to reduce the sucrose concentration. This mixture was then incubated with pre-washed Neutravidin resin (Pierce/Thermo, 29200) overnight at 4°C. The following morning, samples were spun at 1,000 xg to settle the resin, and the depleted supernatant was retained. The resin was washed four times in polysome lysis buffer (as above, without protease inhibitors) for 5 min. Following washes, the resin was resuspended in Buffer RLT and RNase-free EtOH (E7148), and RNA was isolated according to the Qiagen RNeasy Mini kit (74104). The deplete supernatant from the resin was concentrated to 250 µL using centrifugal concentrators (10K cut-off, Amicon, UFC801008). The concentrated supernatant was mixed with buffer RLT and RNase-free EtOH, and RNA was isolated according to the Qiagen RNeasy Mini kit. Biological triplicate samples were sent to Novogene (San Diego, California) where they were library prepped by poly-A purification, reverse transcription, and low-input amplification. Samples were sequenced in a single lane on a NovaSeq X Plus with a minimum of 20 million paired reads (150 bp) per sample. Following sequencing, data underwent Novogene’s data QC and was mapped to the mouse genome.

### Puromycin proximity ligation assay

Cells were treated with 200 µM cycloheximide (Thomas Scientific, C861D14) and 1 mM puromycin (Amresco, J593) for 10 minutes before fixation in -20°C methanol (for DP) or PFA (for all others). For wounding studies, cells were scratch wounded with a razor blade 30 minutes before treatment. After fixation and primary antibody labeling, cells were washed with 3x 5 minutes with buffer A (10 mM Tris, pH 7.4, 150 mM NaCl, 0.05% Tween-20). Secondary antibodies (Sigma Duolink Red Fluorescence Kit, DUO92008) were applied and incubated at 37°C for 1 hour, followed by 3x 5min washes in buffer A. Ligation solution was applied for 30 minutes at 37°C, followed by 2x 5 min washed in buffer A. Amplification solution was next applied and coverslips incubated for 100 min at 37°C. After 3x 5-minute washes, coverslips were stained with Alexa-488 phalloidin (Invitrogen, A12379) and Hoechst, and were sealed and imaged. For quantitation in ImageJ, Z-projections were thresholded, and cortical masks applied (from F-actin signal). Puncta under the mask and in the cytoplasm were determined as well as areas that were covered by the mask. For wounding studies, quantitations were made of puncta within the first two cell layers (for wound) and at least 6 cell layers away (for distal).

### Clear-CLIP

Clear-CLIP was performed as described ^66^, except that Streptavidin beads rather than Ago2 antibody coupled were used. DP-BirA were induced, and biotin labeled as described above. After washing with cold PBS cells were irradiated with 300mJ/cm2 UVC (254nM wavelength), scraped from the plates in cold PBS and pelleted by centrifugation at 1,000g for 2 minutes. Cells were lysed on ice with occasional vortexing in lysis buffer (50 mM Tris-HCl pH 7.4, 100 mM NaCl, 1mM MgCl_2_, 0.1 mM CaCl_2_, 1% NP-40, 0.5% Sodium Deoxycholate, 0.1% SDS) containing 1X protease inhibitors (Roche #88665) and RNaseOUT (Invitrogen #10777019) at 4 µl/ml final concentration. Next, TurboDNase (Invitrogen #AM2238, 10 U), RNase A (0.13 µg), and RNase T1 (0.13 U) were added and incubated at 37°C for 5 minutes with occasional mixing. Samples were placed on ice and then centrifuged at 16,160 g at 4°C for 20 minutes with 25 µl of Streptavidin beads being used per IP. Beads were added to the lysate and rocked for 2 hours at 4°C. All steps after the IP were done on bead until samples were loaded into the polyacrylamide gel. Beads were washed 3 times with cold High Salt Clip Wash Buffer (50 mM Tris-HCl pH 7.4, 1 M NaCl, 1 mM EDTA, 1% NP-40, 0.5% sodium deoxycholate, 0.1% SDS) for 3 minutes with rocking. Samples were then washed 2 times with PNK wash buffer (20 mM Tris-HCl pH 7.4, 10 mM MgCl_2_, 0.2% Tween-20), then phosphorylated at 37°C for 20 minutes in 50 µl of PNK mixture: 41.8 µl H_2_O, 5 µl 10X PNK buffer (NEB), 1 µl RNaseOUT, 1.67 µl ATP (30 mM), 0.5 µl T4 PNK - 3’ phosphatase minus (NEB M0236L). Samples were then washed 3 times. miRNA-mRNA ligation was carried out overnight at room temperature in 49.25 µl H_2_O, 30 µl 50% PEG-8000 (NEB #B1004) 10 µl 10X T4 RNA ligation buffer (NEB), 2.5 µl RNaseOUT, 1 µl ATP (100 mM), 1 µl BSA (10 mg/ml), 6.25 µl T4 RNA ligase 1 (10 U/ml – NEB M0204). The following morning, an additional 2.5 µl T4 RNA ligase 1 (10 U/ml) and 1 µl ATP (100 mM) were added, and ligation was continued for another 5 hours. Samples were then washed twice with lysis buffer, once with PNK/EDTA/EGTA (50 mM Tris pH 7.4, 10 mM EDTA, 10 mM EGTA, 0.5% Igepal) and twice more with PNK wash buffer. Next, samples were treated with phosphatase at 37°C for 20 minutes with 41 µl H_2_O, 5 µl 10X FastAP buffer, 3 µl FastAP enzyme (Thermo Fisher #EF0651) and 1 µl RNaseOUT. Samples were then washed two times with PNK wash buffer, and 3’ adapter ligation was performed on beads overnight. Samples were then washed twice with PNK buffer. Samples were then heated to 70°C for 10 minutes with occasional agitation and supernatant was separated from beads. RNA was isolated by following the protocol published for irClip (Zarnegar et al. 2016): 15 µl of Proteinase K (Thermo scientific EO0491) at 20 mg/ml was added to 285 µl of Proteinase K/SDS buffer (100 mM TrisHCl pH 7.5, 50 mM NaCl, 1 mM EDTA, 0.2% SDS). This solution was heated to 37°C for 20 minutes to inactivate any RNases and incubated at 50C for 1 hour.

Samples were briefly centrifuged and then 375 µl of saturated phenol/chloroform/isoamyl alcohol (25:24:1) was added and incubated for 10 minutes at 37°C. Samples were then centrifuged at 16,000g at room temperature for 3 minutes, the aqueous layer was removed to a new tube and precipitated overnight at -20°C with 2 µl Glycoblue (ThermoFisher #AM9516) and 900 µl of 100% ethanol. RNA was pelleted by centrifugation at 16,160g at 4°C for 20 minutes and the supernatant removed. The pellet was washed with 70% ethanol and left to air dry at room temperature for 5 minutes. The 5’ adapters were ligated by adding 8 µl of mix to the pellet (2 µl 50% PEG-8000, 1µl 10X NEB RNA ligation buffer, 1 µl 10mM ATP, 4 µl H_2_O). Samples were heated briefly to 95°C, placed on ice and additional components were added: 0.5 µl RNaseOUT, 1 µl T4 RNA ligase 1 (10U/µl – NEB M0204), 0.5 µl 100 µM 5’ RNA linker (Blocked at 5’ end & NNNN at 3’ end for bias reduction and barcoding). Ligation was carried out at 37°C for 4 hours with rocking. RT-PCR was then carried out in ligation buffer by adding 8.5 µl of the following mix to the sample: 4 µl 5X first-strand buffer (Invitrogen), 1.5 µl 100mM DTT, 2 µl 1 mM RT primer, 1 µl 10mM dNTPs. Samples were heated to 65°C for 5 minutes, transferred to a PCR tube and then enzymes were added: 1 µl Superscript III (Thermo Fisher #18080093) and 0.5 µl RNaseOUT. RT reaction was then performed in a thermocycler: 50°C for 1 hour, 85°C for 5 minutes, and then hold at 4°C. Libraries were then amplified by PCR. Products were run on a 9% acrylamide gel and then stained with SYBR gold (ThermoFisher #211494) at 1:10,000 for 10 minutes. The appropriate gel region was excised, frozen at -80°C, resuspended in HSCB buffer overnight. The next day the gel slurry was transferred to a 0.22 µm filter tube and spun at 16,000g for 20 minutes at room temperature. Samples were precipitated overnight and then pelleted at 16,160g at 4°C for 20 minutes. The pellet was washed with 70% ethanol, air dried for 5 minutes and then resuspended in 20 µl of H_2_O. Sequencing barcodes were added by PCR. Products were run on a 9% acrylamide gel and stained with SYBR gold as above. Products corresponding to sizes approximately 144-200 base pairs were excised and isolated from the gel. Libraries were mixed in equal amounts and sequenced on an Illumina HiSeq 4000 by the Microarray and Genomics Core at the University of Colorado Anschutz Medical Campus.

## Plasmids Used/Generated

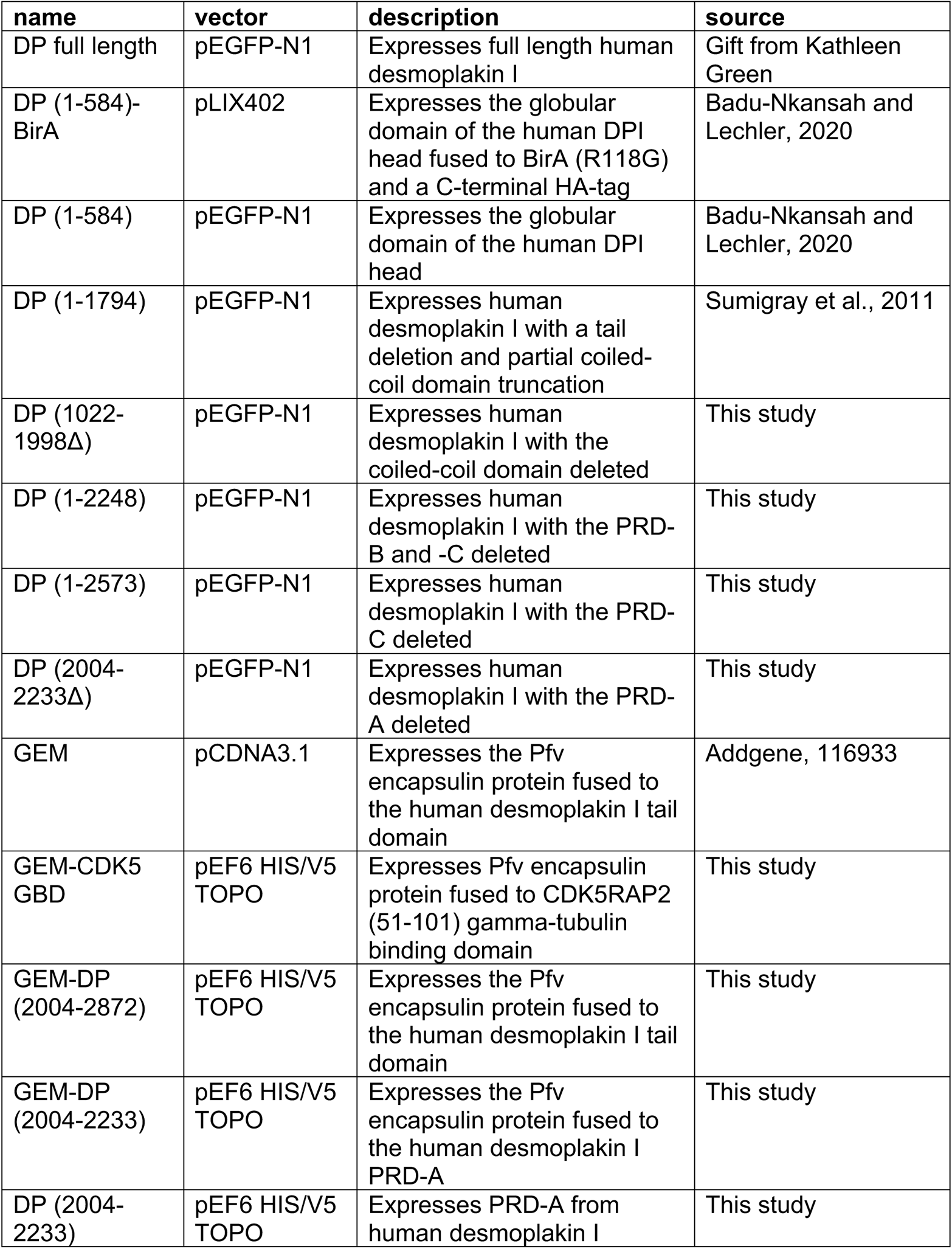

## Data Availability

RNA-Seq and CLEAR-CLIP data have been deposited in the Gene Expression Omnibus under accession codes GSE281632 and GSE283064. Any other data supporting the findings of this study are available from the corresponding author upon request.

## Acknowledgements

We thank Julie Underwood for lab management and care of the mice used in this study. This work was supported by grants R01-AR067203, R01-AR081081 and R01-AR083352 to TL; R01-GM139480 to CVN; and R01AR066703 to RY.

---

**Figure S1:**
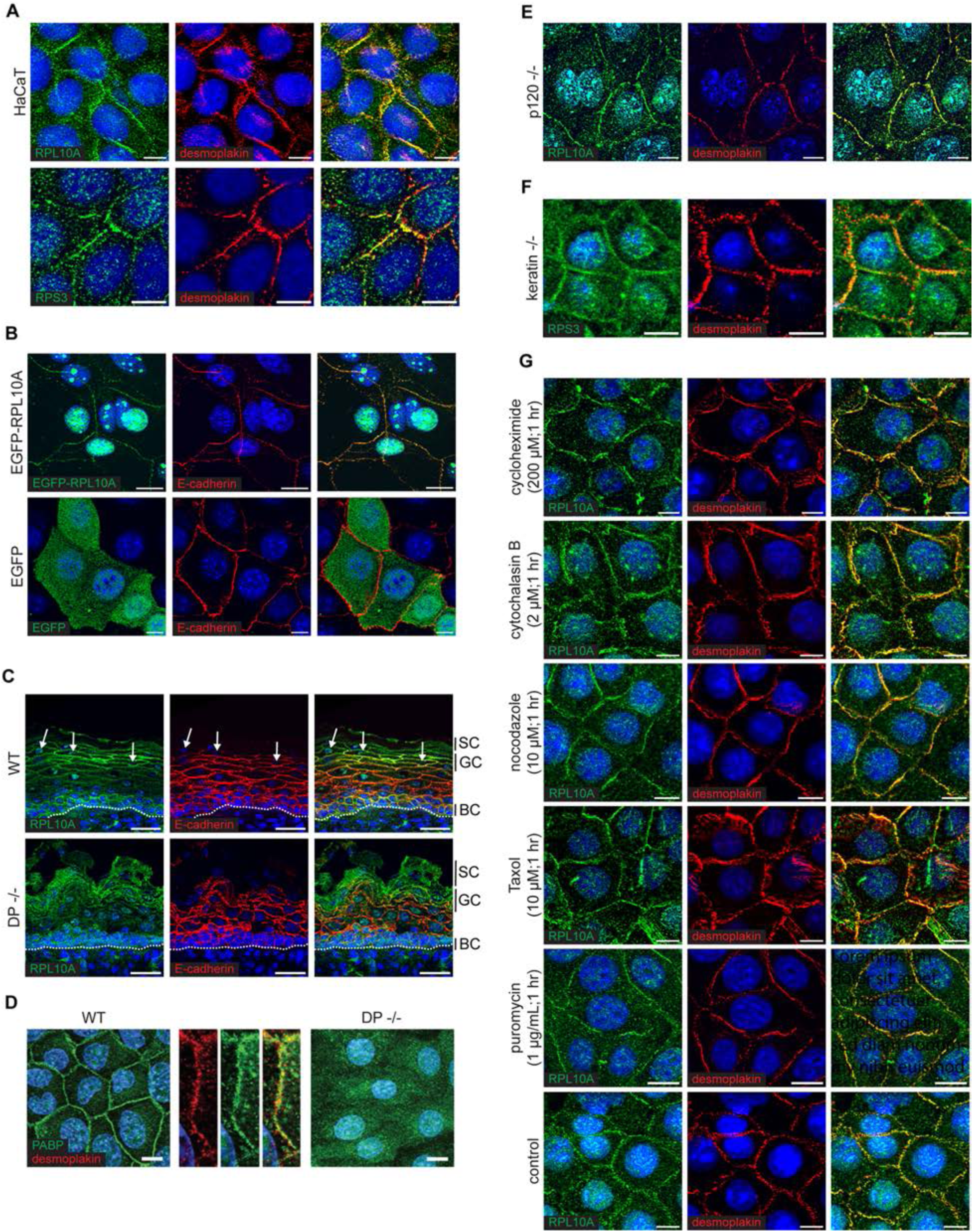
Mechanisms of cortical ribosome localization. (A) Immunofluorescence of RPL10A or RPS3 and desmoplakin in HaCaT human keratinocytes (scale bar = 10 µm). (B) EGFP-RPL10A or control EGFP expressing mouse keratinocytes counterstained with E-cadherin (scale bar = 10 µm). (C) Immunofluorescence of RPL10A and E-cadherin in wild-type and desmoplakin-null mouse E18.5 epidermis. Dotted line represents basement membrane (BC = basal cell, GC = granular cell, SC = stratum corneum; scale bar = 30 µm). (D) Immunofluorescence of poly-A binding protein (PABP) in wild-type and desmoplakin-null mouse keratinocytes. Inset shows close-up of cell-cell border in wild-type cells with desmoplakin counterstain (scale bar = 10 µm). (E) Immunofluorescence of RPL10A and desmoplakin in p120 catenin-null mouse keratinocytes (scale bar = 10 µm). (F) Immunofluorescence of RPS3 and desmoplakin in keratin-null mouse keratinocytes (scale bar = 10 µm). (G) Immunofluorescence of RPL10A and desmoplakin in wild-type mouse keratinocytes treated with cycloheximide, puromycin, cytochalasin B, nocodazole, paclitaxel, or vehicle control at the indicated times and concentrations.

**Figure S2:**
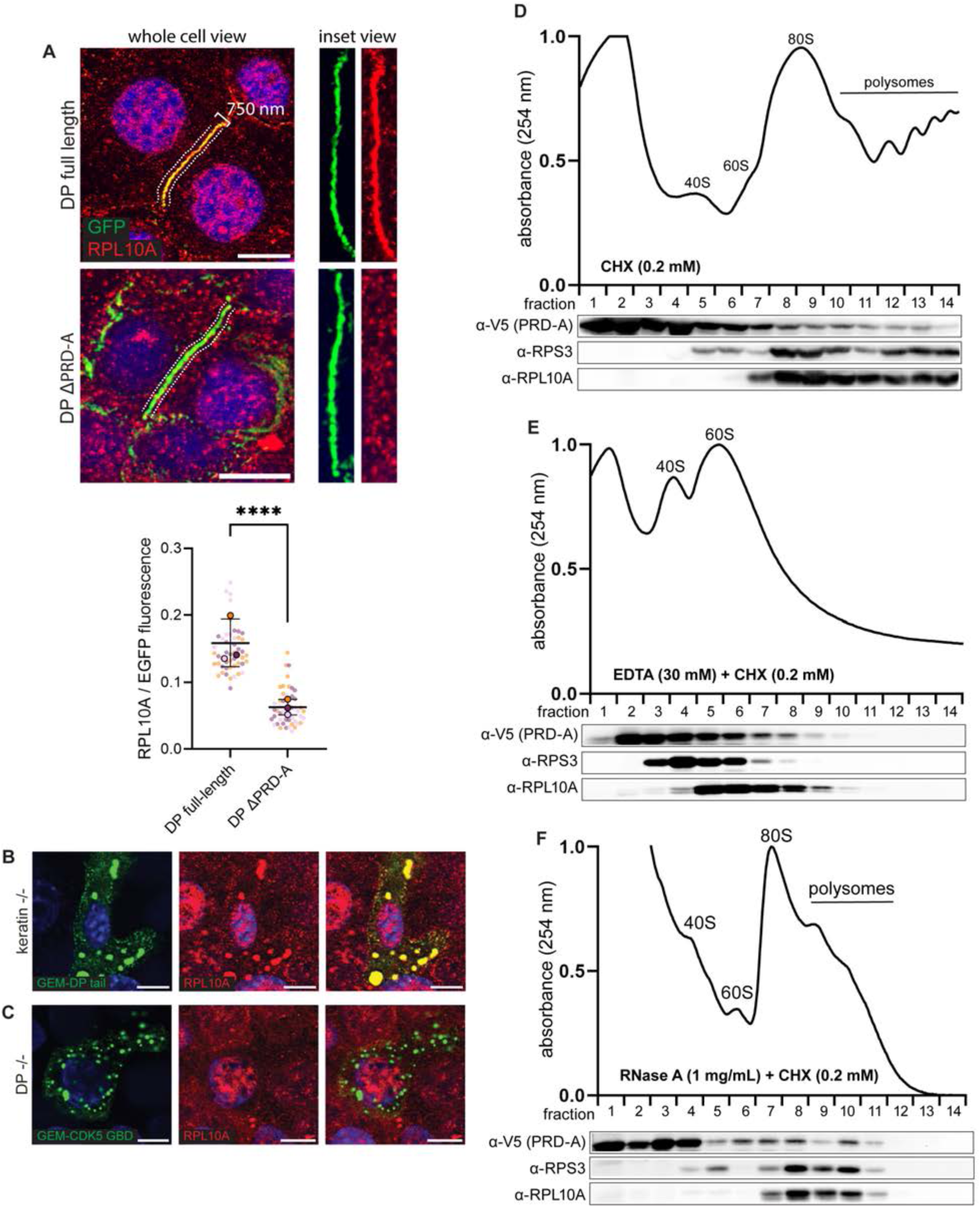
The PRD-A domain of desmoplakin co-sediments with ribosomes and is necessary for cortical localization. (A) Example images of desmoplakin-null cells transfected with full-length or ΔPRD-A desmoplakin constructs with close-up view of the cell-cell boundary. Accompanying quantification of image data measuring the ratio of average RPL10A to EGFP fluorescence intensity across 750 nm-wide cell-cell boundaries using fitted lines (as illustrated) in desmoplakin-null keratinocytes transfected with full-length or ΔPRD-A desmoplakin constructs (full-length n = 51 cells, ΔPRD n = 51 cells; 3 biological replicates; p < 0.0001). (B) Keratin-null mouse keratinocytes transfected with GEM-DP tail and co-stained for RPL10A (scale bar = 10 µm). (C) desmoplakin-null mouse keratinocytes transfected with GEM-Cdk5 GBD (a GEM fusion known to aggregate) and co-stained for RPL10A (scale bar = 10 µm). (D-F) Polysome profile and fractions from cells expressing soluble V5-tagged desmoplakin PRD-A and lysed in (D) normal conditions (only 0.2 µM CHX) (E) 30 mM EDTA or (F) RNase A (1 mg/mL; 30 min). Proteins from fractions were separated by SDS-PAGE and blotted for V5 (PRD-A), RPS3, and RPL10A. Higher fraction numbers represent higher sucrose density.

**Figure S3:**
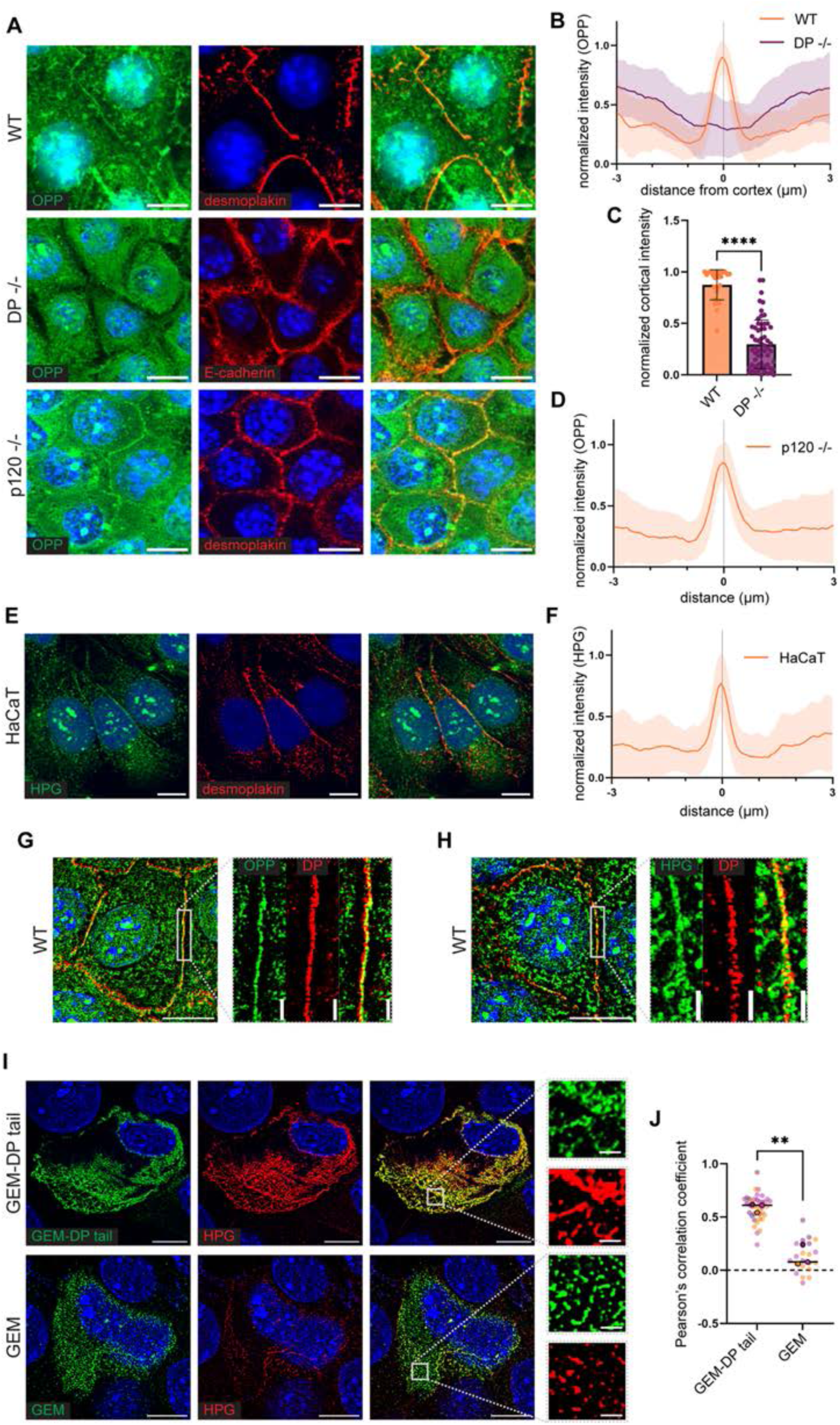
Evidence for local translation at the cell cortex. (A) Click-chemistry labeling of O-propargyl-puromycin (OPP; 5 min pulse) and immunofluorescence of desmoplakin or E-cadherin in wild-type, desmoplakin-null, or p120-null mouse keratinocytes (scale bar = 10 µm). (B) Line scans of OPP fluorescence intensity measured across the cell cortex of wild-type and desmoplakin-null keratinocytes. Shaded area represents the standard deviation (WT n = 31 cells, DP-/- n = 60 cells; 3 biological replicates). (C) Relative intensity of OPP fluorescence at the center of the cell-cell boundary (WT n = 31 cells, DP-/- n = 60 cells; 3 biological replicates; p < 0.0001). (D) Line scans of OPP fluorescence intensity measured across the cell cortex in p120-null keratinocytes. Shaded area represents the standard deviation (n = 51 cells; 3 biological replicates). (E) Click-chemistry labeling of HPG (5 min pulse) and immunofluorescence of desmoplakin in HaCaT human keratinocytes (scale bar = 10 µm). (F) Line scans of HPG fluorescence intensity measured across the cell cortex in HaCaT human keratinocytes. Shaded area represents the standard deviation (n = 40 cells; 3 biological replicates). (G) Lattice-SIM close-up images of OPP labeling at the cell-cell boundary in wild-type mouse keratinocytes (SIM scale bar = 0.5 µm). (G) Lattice-SIM close-up images of HPG labeling at the cell-cell boundary in wild-type mouse keratinocytes (SIM scale bar = 0.5 µm). (I) Lattice-SIM images of click-chemistry labelled HPG (3 min pulse) and immunofluorescence of desmoplakin-null cells transfected with GEM-DP tail or cytoplasmic GEM plasmids (SIM scale bar = 0.5 µm). (J) Pearson’s correlation coefficient (PCC) of EGFP and HPG fluorescence from cells transfected as in (I) (GEM-DP tail n = 38 cells, GEM n = 22 cells; 3 biological replicates; p < 0.0018).

**Figure S4:**
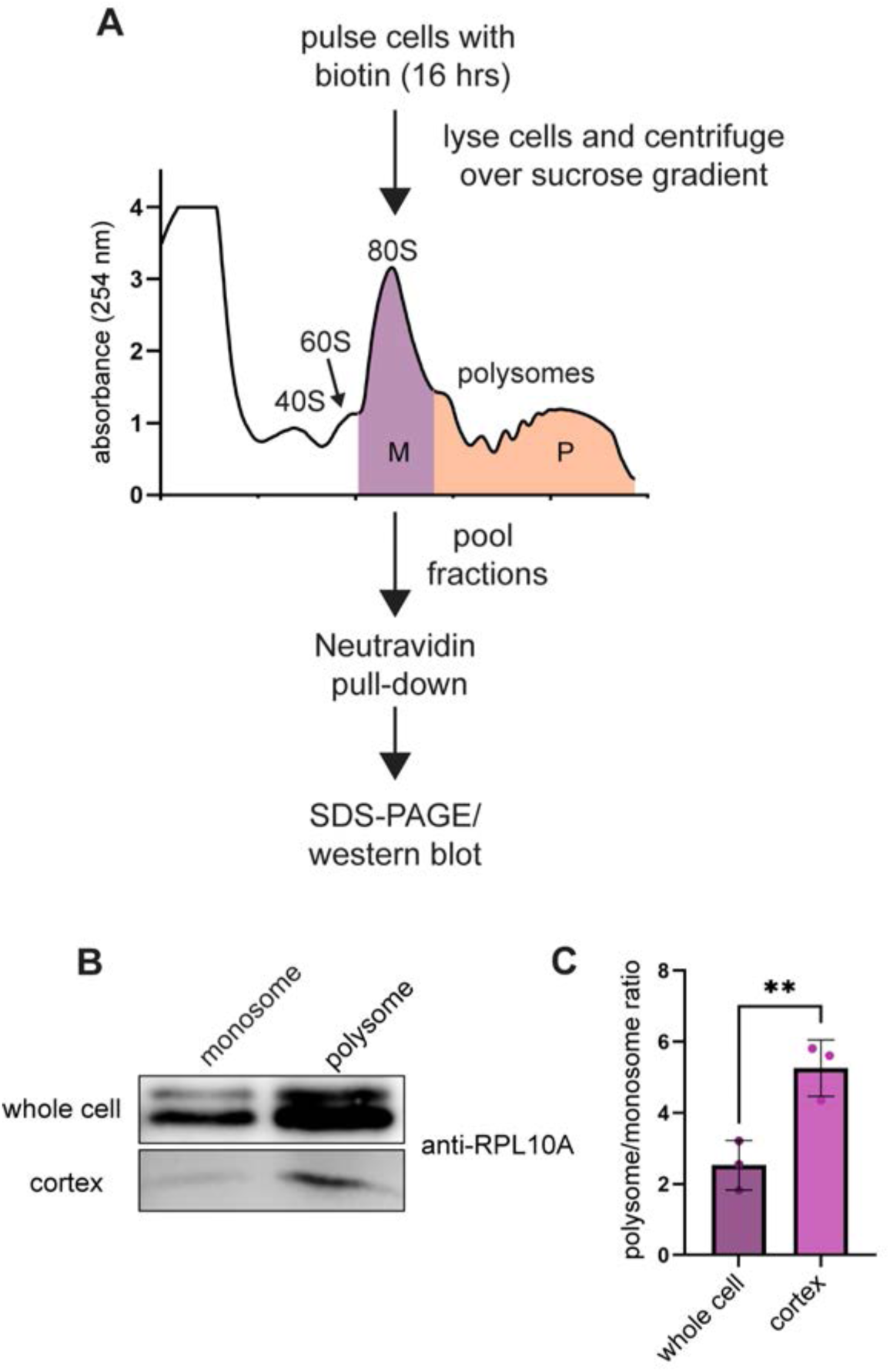
Cortical ribosomes are polysome-associated. (A) Schematic of cortical ribosome labeling, purification, and monosome/polysome isolation. (B) Western blot of cortically isolated and whole cell RPL10A from pooled monosome and polysome fractions. (C) Ratio of polysome to monosome band intensities from RPL10A western blots (n = 3 biological replicates; p = 0.0037).

**Fig. S5.**
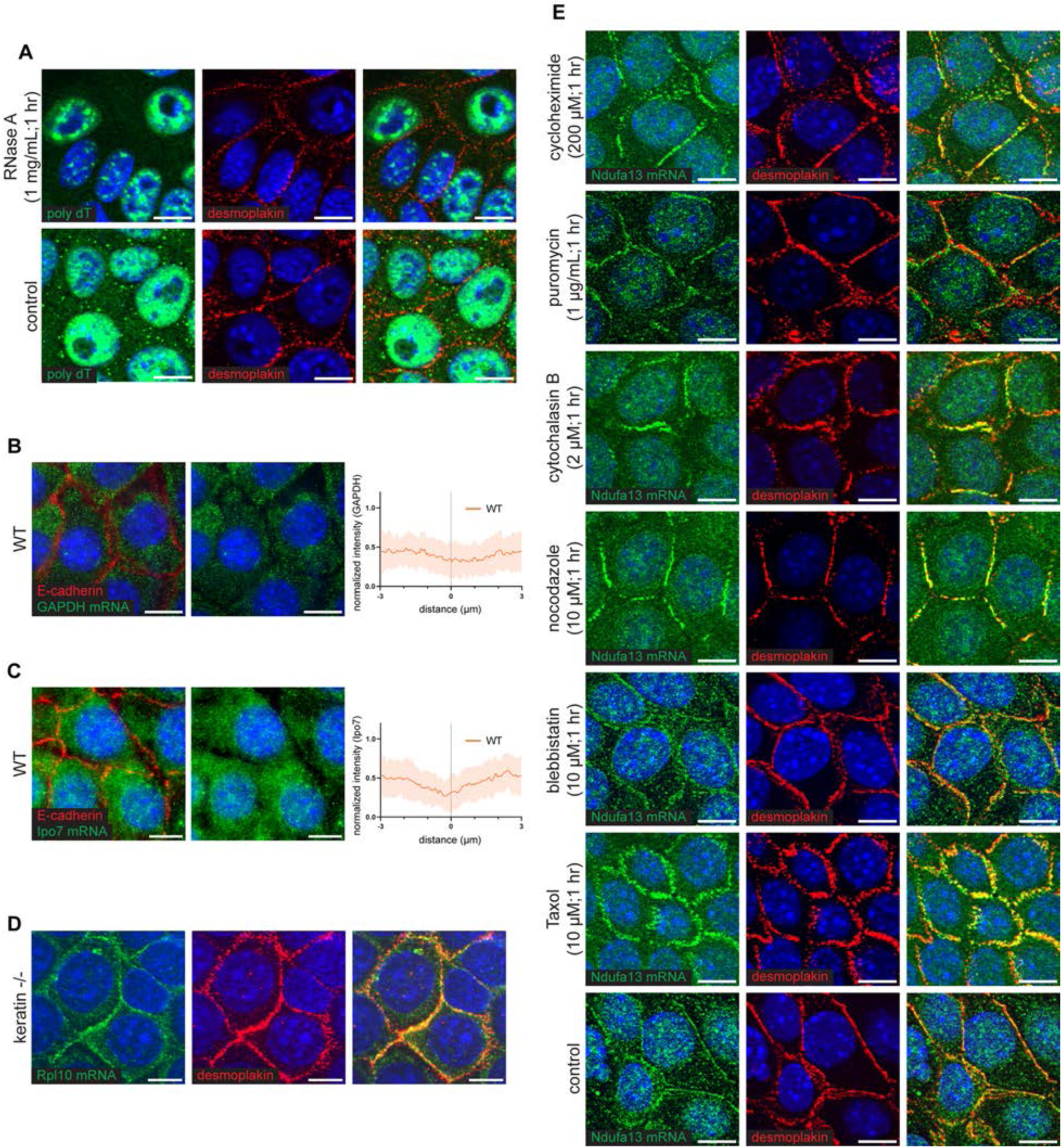
Mechanisms of Ndufa13 mRNA localization. (A) Poly-dT FISH in wild-type mouse keratinocytes treated or untreated with RNase A (1 mg/mL, 1 hr) and co-stained for desmoplakin (scale bar = 10 µm). Same day control with intensity scaling matched. (B) smFISH of GAPDH mRNA and E-cadherin immunofluorescence in wild-type mouse keratinocytes and line scans of GAPDH probe intensity measured across the cell cortex. (C) smFISH of Ipo7 mRNA and E-cadherin immunofluorescence in wild-type mouse keratinocytes and line scans of Ipo7 probe intensity measured across the cell cortex. (D) smFISH labeling of RPL10 mRNA and desmoplakin immunofluorescence in keratin-null mouse keratinocytes (scale bar = 10 µm). (E) smFISH of Ndufa13 mRNA and desmoplakin immunofluorescence in wild-type mouse keratinocytes treated with cycloheximide, puromycin, cytochalasin B, nocodazole, blebbistatin, paclitaxel, or vehicle control at the indicated times and concentrations.

**Figure S6:**
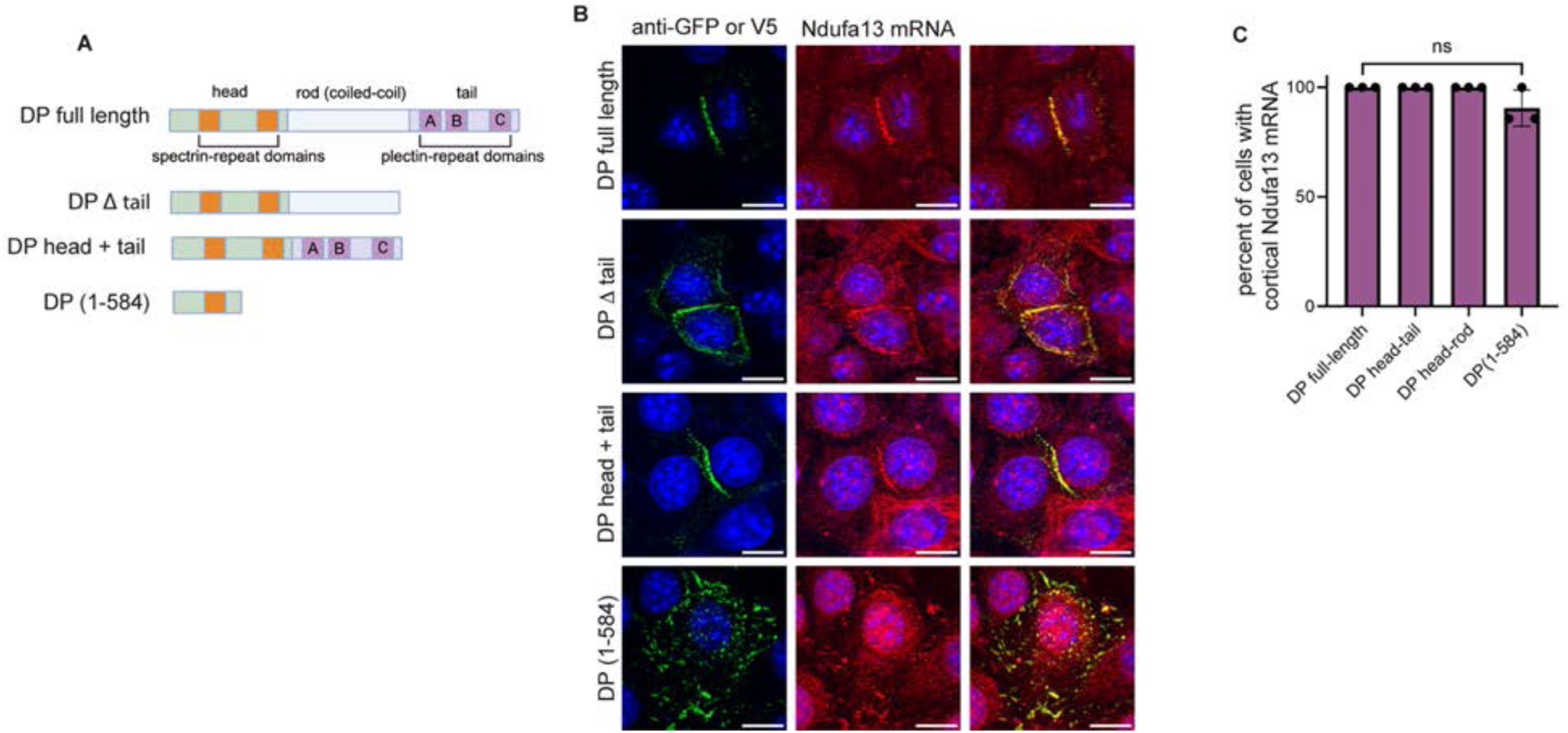
Mechanisms of mRNA cortical localization. (A) Overview of desmoplakin constructs tested for their ability to rescue Ndufa13 mRNA localization. (B) Desmoplakin-null cells transfected with various GFP or V5-tagged desmoplakin constructs, stained for GFP, and probed for Ndufa13 mRNA. (C) Percent of desmoplakin-null cells transfected with various desmoplakin constructs that show rescued cortical Ndufa13 hybridization (n = >10 cells from 3 biological replicates; p < 0.06).

**Figure S7:**
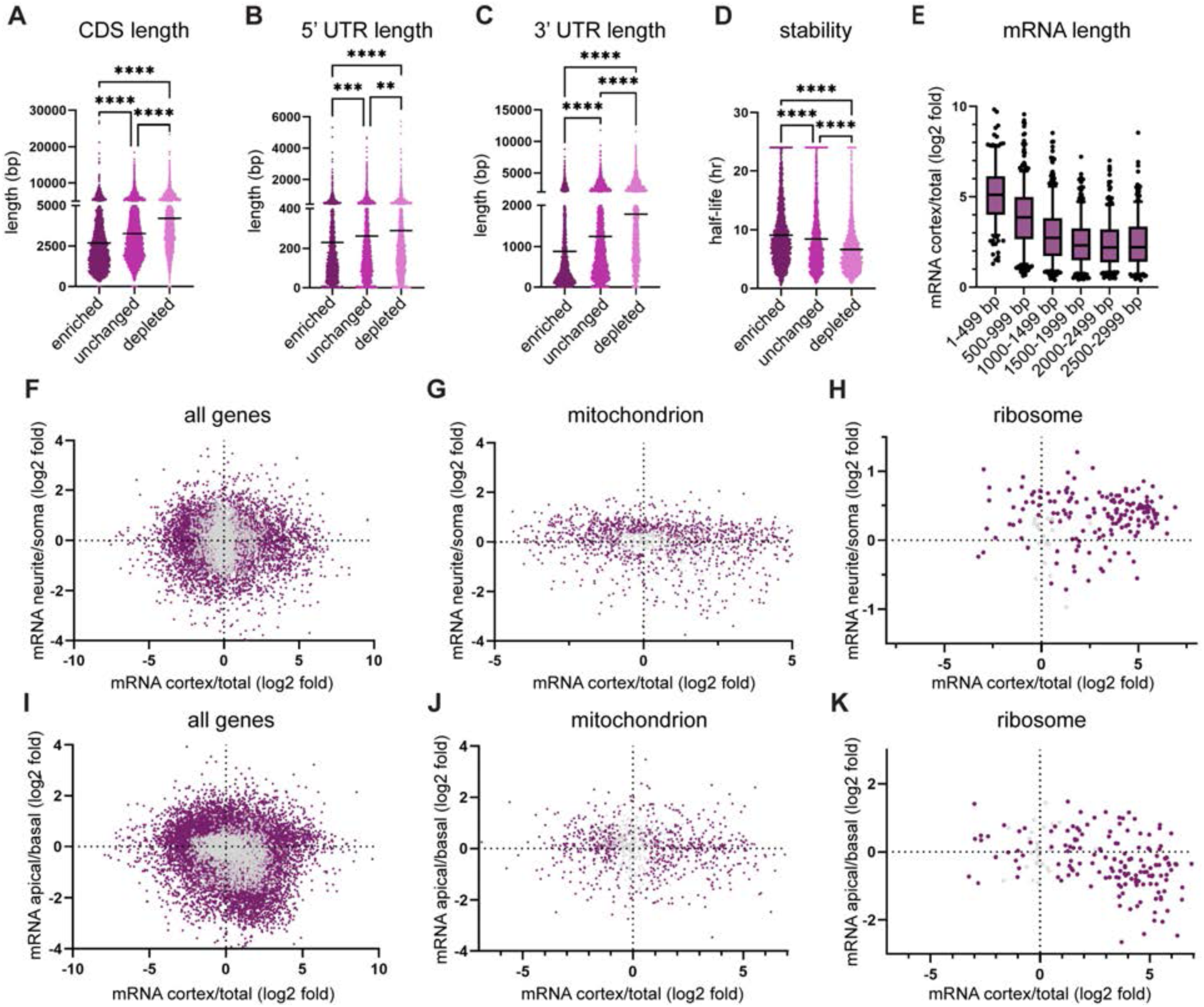
mRNA length and stability are correlated with localization and mRNA localization is regulated differently between cell types. (A) Coding-sequence length of mRNAs that were significantly enriched (Log2FC > 0.15, p < 0.05), unchanged (Log2FC -0.15 – 0.15, p > 0.05), or depleted (Log2FC > -0.15, p < 0.05) (enriched n=2489, unchanged n=4280, depleted n=2595; p < 0.0001). (B) 5’ UTR sequence length of mRNAs that were significantly enriched, unchanged, or depleted (p < 0.0001). (C) 3’ UTR sequence length of mRNAs that were significantly enriched, unchanged, or depleted (p < 0.0001). (D) Stability of mRNAs that were significantly enriched, unchanged, or depleted (p < 0.0001). (E) Cortical enrichment of mRNAs binned by sequence length (1-499 bp n = 290, 500-999 bp n = 578, 1000-1499 bp n = 560, 1500-1999 bp n = 461, 2000-2499 bp n = 436, 2500-2999 bp n = 373, p < 0.0001). (F) mRNA localization at the cortex and within the neurite. Grey: insignificant and Purple: significant (PCC = -0.15, n = 9944, p < 0.0001). (G) Mitochondrion-related mRNA localization at the cortex and within the neurite (PCC = -0.15, n = 1148, p < 0.0001). (H) Ribosome-related mRNA localization at the cortex and within the neurite (PCC = 0.06, n = 170, p = 0.47). (I) mRNA localization at the cortex and within the enterocyte. Grey: insignificant and Purple: significant (PCC = -0.04, n = 5073, p = 0.004). (J) Mitochondrion-related mRNA localization at the cortex and within the neurite (PCC = -0.08, n = 711, p = 0.041). (K) Ribosome-related mRNA localization at the cortex and within the neurite (PCC = -0.385, n = 149, p < 0.0001).

**Figure S8:**
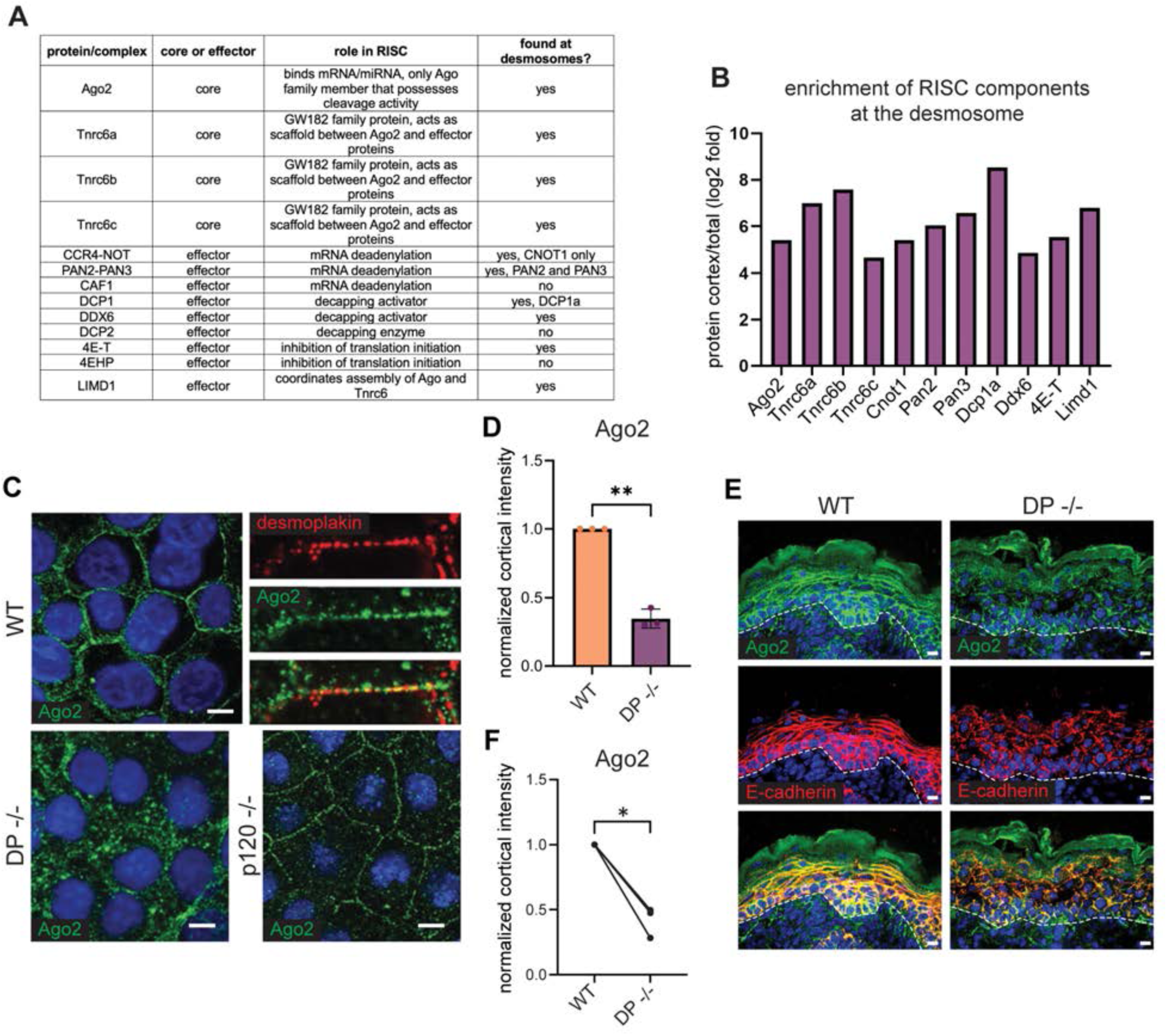
Validation of desmosome dependent RISC cortical localization. (A) Table containing descriptions of RISC-associated proteins and their association with the desmosomal proteome. (B) Desmosomal enrichment of RISC components. (C) Representative immunofluorescence images of Ago2 in wild-type, desmoplakin-null, and p120-null mouse keratinocytes. Wild-type cells show at inset counterstained with desmoplakin (scale bar = 10 μm). (D) Relative intensity of Ago2 at the center of the cell-cell boundary for wild-type, desmoplakin-null, and p120-null mouse keratinocytes (n = 3 biological replicates; WT vs. DP-/- p = 0.0067; WT vs. p120-/- p = 0.0155; DP vs. p120-/- p = 0.1232). (E) Staining for Ago2 and E-cadherin in back skin of E18.5 wild-type and desmoplakin-null mouse embryos. Basement membrane indicated by a white dotted line (scale bar = 10 μm). (F) Relative intensity of Ago2 at the center of the cell-cell boundary for WT and desmoplakin-null tissue samples from (E) (n = 3 mice).

**Figure S9:**
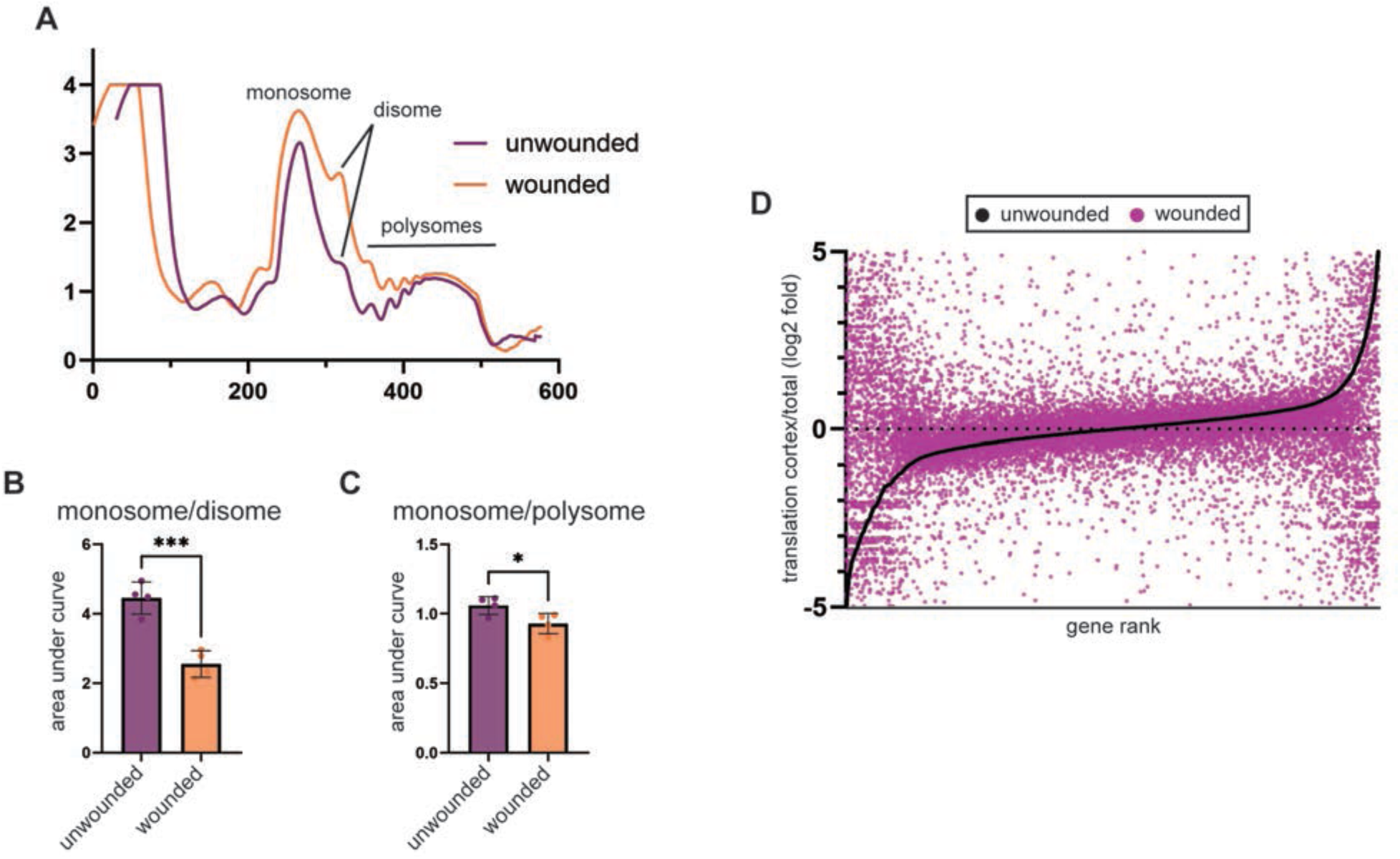
Perturbing cell-cell adhesion affects translation in keratinocytes. (A) Polysome traces of wounded and unwounded mouse keratinocytes collected at 16 hrs post wounding. (B) Monosome/disome area under curve ratios calculated from wounded and unwounded polysome traces (n = 4 biological replicates, p = 0.0007). (C) Monosome/polysome area under curve ratios calculated from wounded and unwounded polysome traces (n = 4 biological replicates, p = 0.0369). (D) Genes listed by rank from least to most cortically translated in unwounded cells (black) and the cortical enrichment of the same genes in the wounded (purple).

**Figure S10.**
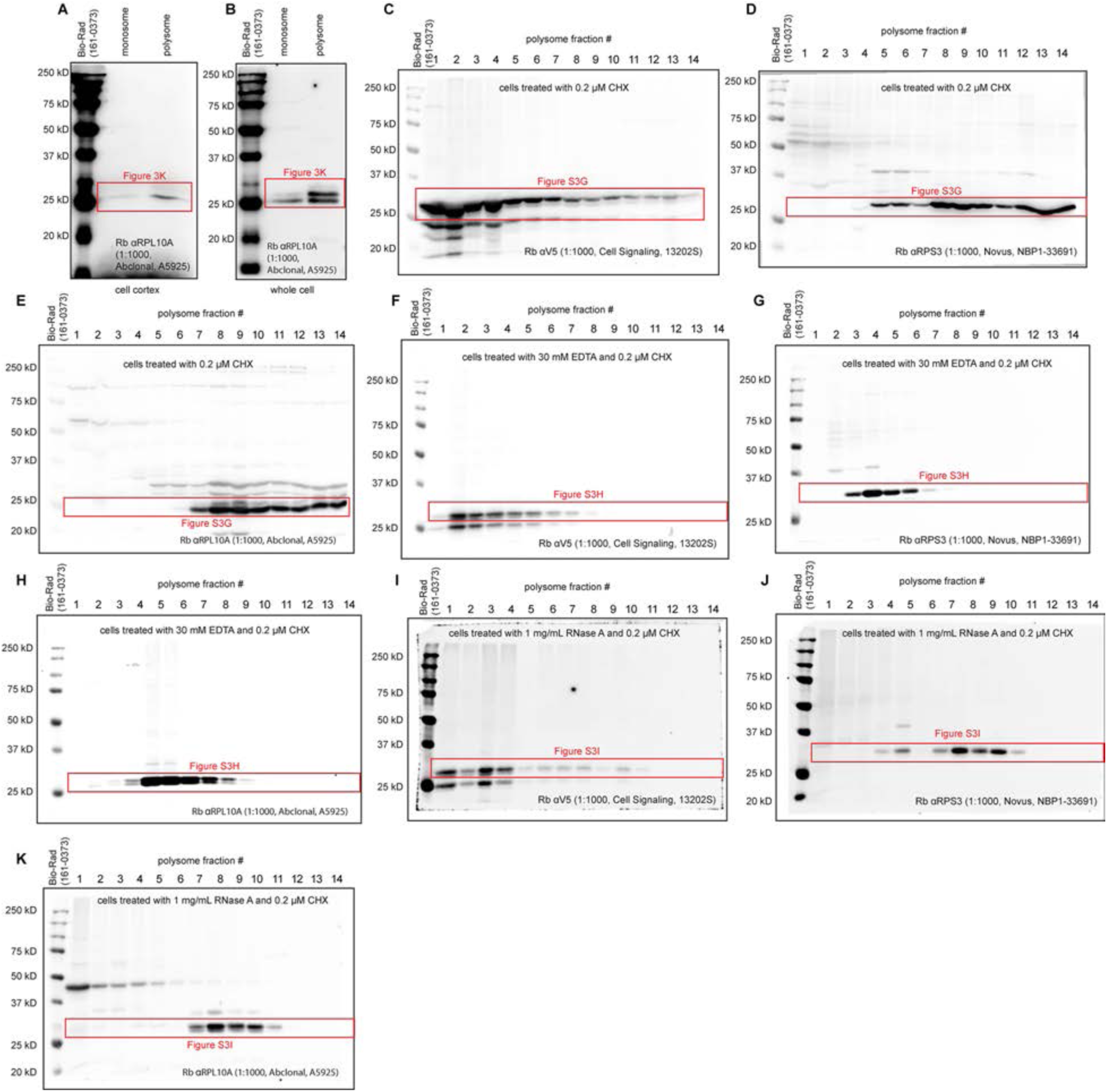
Whole western blot images of data from Figures 3 and S2. (A) Cortical and (B) whole cell monosome/polysome western blot data. (C-E) Full western blot data of cell lysate treated with 0.2 µM cycloheximide then centrifuged over a sucrose gradient (15%-50%) to fractionate. (F-H) Full western blot data of cell lysate treated with 30 mM EDTA and 0.2 µM cycloheximide then centrifuged over a sucrose gradient (15%-50%) to fractionate. (I-K) Full western blot data of cell lysate treated with 1 mg/mL RNase A and 0.2 µM cycloheximide, centrifuged over a sucrose gradient (15%-50%) to fractionate. Higher fraction numbers represent higher sucrose density.

